# Glutamine metabolism inhibition has dual immunomodulatory and antibacterial activities against *Mycobacterium tuberculosis*

**DOI:** 10.1101/2023.02.23.529704

**Authors:** Sadiya Parveen, Jessica Shen, Shichun Lun, Liang Zhao, Benjamin Koleske, Robert D. Leone, Rana Rais, Jonathan D. Powell, John R. Murphy, Barbara S. Slusher, William R. Bishai

**Author notes:** Corresponding author William R. Bishai, Johns Hopkins School of Medicine, 1550 Orleans St, CRB2 Rm 108, Baltimore, MD 21287.

## Abstract

As one of the most successful human pathogens, *Mycobacterium tuberculosis* (*Mtb*) has evolved a diverse array of determinants to subvert host immunity and alter host metabolic patterns. However, the mechanisms of pathogen interference with host metabolism remain poorly understood. Here we show that a novel glutamine metabolism antagonist, JHU083, inhibits *Mtb* proliferation in vitro and in vivo. JHU083-treated mice exhibit weight gain, improved survival, a 2.5 log lower lung bacillary burden at 35 days post-infection, and reduced lung pathology. JHU083 treatment also initiates earlier T-cell recruitment, increased proinflammatory myeloid cell infiltration, and a reduced frequency of immunosuppressive myeloid cells when compared to uninfected and rifampin-treated controls. Metabolomics analysis of lungs from JHU083-treated *Mtb*-infected mice revealed reduced glutamine levels, citrulline accumulation suggesting elevated NOS activity, and lowered levels of quinolinic acid which is derived from the immunosuppressive metabolite kynurenine. When tested in an immunocompromised mouse model of *Mtb* infection, JHU083 lost its therapeutic efficacy suggesting the drug’s host-directed effects are likely to be predominant. Collectively, these data reveal that JHU083-mediated glutamine metabolism inhibition results in dual antibacterial and host-directed activity against tuberculosis.

## INTRODUCTION

Tuberculosis (TB) remains a leading infectious disease killer causing ∼1.5 million deaths worldwide in 2020 alone ^1^. The causative bacterium, *Mycobacterium tuberculosis* (*Mtb*), is one of the most successful global pathogens, and it has evolved multiple mechanisms to evade and suppress the host immune system ^2^. While effector CD4^+^ T-cells are considered one of the most potent factors in host containment of the pathogen, individuals with active TB disease fail to mount adequate cellular immune responses ^3,4^, and the mechanisms by which *Mtb* infection leads to failed effector T-cell responses remain poorly understood.

Suppression of effector T-cell immunity is also one of the hallmarks of the tumor microenvironment ^5^. Recently, a novel glutamine (Gln) metabolism antagonist drug, JHU083, has been shown to reprogram host immunometabolic signatures and improve effector T-cell immunity in several murine tumor models ^6-8^. High Gln metabolism in tumors were found to be closely associated with an abundance of immunosuppressive cells such as myeloid-derived suppressor cells (MDSCs) and T regulatory cells (Tregs) with a concomitant inhibition of effector T cells^9,10^. JHU083 administration not only reduced Gln utilization but also enhanced the antitumor activity of effector T cells ^6,7^. JHU083 and a closely related drug Sirpiglenastat, which is in human clinical trials, are prodrugs of the glutamine antagonist, 6-diazo-5-oxo-L-norleucine (DON), which has shown to have anticancer activity in preclinical models and early clinical trials ^8,11^. DON irreversibly inhibits glutamine-utilizing enzymes including glutaminases, glutamine synthetases as well as multiple glutamine amidotransferases involved in the biosynthesis of purines, pyrimidine, coenzymes, hexosamines and amino acids^12^. While DON showed robust antitumor activity in preclinical and early clinical trials, its development was hampered by dose-limiting toxicities which were mainly gastrointestinal (GI) track-related^12^. To circumvent these GI toxicities and enhance its therapeutic index, prodrugs of DON were designed to preferentially deliver DON to tumors while minimizing release in the gut^11,13^. One such prodrug, JHU083, is orally available and is converted to DON by ubiquitous peptidases and esterases, has fewer side effects, and has shown the ability to enhance T-cell immunity in the tumor microenvironment^8,11^. Since blunted effector T-cell immunity are characteristics of both the cancer microenvironment and the tuberculous granuloma ^14^, we hypothesized that JHU083 might also enhance effector T-cell immunity by a similar immunometabolic reprogramming mechanism and thereby improve host containment of TB disease progression.

In support of this hypothesis, early seminal work by Horwitz and colleagues indicated that inhibition of bacterial glutamine (Gln) metabolism may have direct antibacterial activity^15-17^. This group showed that *Mtb* possesses four glutamine synthetase enzymes. Among these four proteins, only GlnA1 is secreted, and was found to be essential for *Mtb* survival in vitro and in vivo^17^. In addition, Horwitz et al. also showed that methionine sulfoximine (MSO), an irreversible inhibitor of glutamine synthetase enzymes, has direct anti-mycobacterial activity with an *Mtb* MIC of 50 μM on solid medium^15^. MSO treatment also reduced the bacillary burden in the guinea pig TB model^15^. These observations, together with the aforementioned immunometabolic reprogramming by a glutamine metabolism antagonist, suggested to us that *Mtb* may employ its extracellular GlnA1 enzyme as a virulence factor to increase Gln metabolism in the granuloma microenvironment, thereby creating an immunosuppressive cellular milieu as has been shown in tumors. We, therefore, hypothesized that JHU083 may serve as a host-directed therapy against TB disease by reprogramming Gln metabolism in granulomas. In light of the known direct antibacterial effects of the glutamine synthetase inhibitor MSO, we considered the possibility that JHU083 might possess direct antibacterial activity as well.

In the present study, we demonstrate that JHU083 has dual immunomodulatory and antibacterial activity against TB disease. In vitro, we show that DON exhibited direct anti-mycobacterial activity against *Mtb* in broth cultures as well as infected bone marrow-derived macrophages (BMDM). Given its better therapeutic index, we evaluated the DON prodrug JHU083 in the murine model of TB. We found that JHU083 administration reduced lung bacillary burden and improved lung pathology, which resulted in weight gain and prolonged survival of the mice. JHU083 administration did not offer any therapeutic benefit in immunocompromised mice infected with *Mtb*. Using rifampin (RIF) monotherapy as a control antibacterial agent, multicolor flow cytometry revealed that JHU083 treatment initiated earlier recruitment of activated T-cells exhibiting follicular helper and naive signatures than did RIF. The increase in the follicular helper T-cell signature was concomitant with increased frequency of activated mature B-cells. These effects were not limited to the lymphoid compartment, as JHU083 administration also reprogrammed myeloid cells towards proinflammatory phenotypes to a greater degree than RIF. JHU083 administration also led to reduced levels of glutamine, accumulation of citrulline which is consistent with higher host-protective NOS activity, and depletion of quinolinic acid, a byproduct of the immunosuppressive metabolite kynurenine. Taken in total, these findings reveal that glutamine antagonism may be beneficial in host *Mtb* containment through a combination of immunomodulatory reprogramming and direct antibacterial killing.

## RESULTS

### 1. DON exhibits modest antibacterial activity against Mtb

Based on the earlier work of Horwitz and colleagues showing the essentiality of GlnA1, the major *Mtb* glutamine synthetase, we hypothesized that JHU083, a glutamine synthetase inhibitor, may also inhibit *Mtb* growth^18^. Using the Alamar blue assay, we tested the anti-mycobacterial activity of JHU083, DON, and MSO, a known inhibitor of *Mtb* GlnA1^15,19^(**Fig 1A and 1B)**. Both JHU083 and DON inhibited *Mtb* H37Rv growth in vitro with an MICs of 1 μg/ml which was four times more potent than MSO (MIC = 4 μg/ml) (**Fig 1C**). RIF was used as the positive control in these assays (MIC = 0.25 μg/ml) (**Fig 1C**).

**Fig 1.**
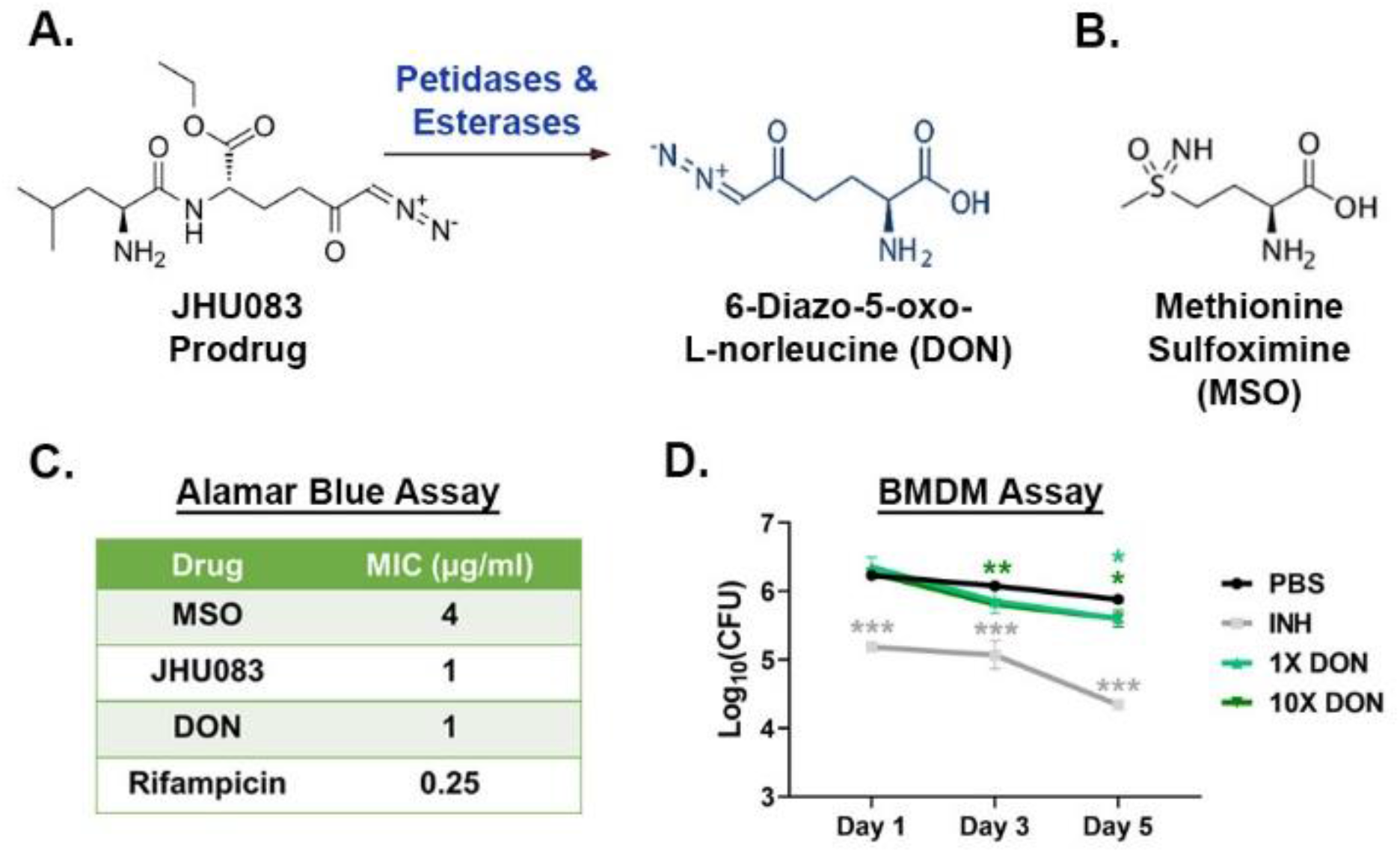
DON has direct anti-mycobacterial activity *in vitro*. (**A)** Chemical structures of the prodrug JHU083 and the active drug DON. Esterases and peptidases ubiquitously present in the blood and tissues convert JHU083 into DON. (**B)** Chemical structure of MSO. (**C)** Table showing the MIC values of these drugs against *Mtb* H37Rv strain determined using Alamar Blue Assay. (**D)** IFNƳ-activated BMDM were infected with *Mtb* H37Rv at an MOI of 2. They were then treated with 1X and 10X MIC concentrations of DON. Isoniazid (INH) was used as the positive control. The cells were lysed at indicated time points and plated on 7H11 selection plates. The assay was performed as described in the “Methods”. Data were plotted as Mean ±SEM. Statistical significance was calculated using a two-tailed student t-test considering unequal distribution. *<0.05, **<0.01, ***<0.001. All the experiments were performed in triplicates.

We also determined the antibacterial activity of DON ex vivo in *Mtb* H37Rv infected murine bone-marrow derived macrophages. DON at 1X and 10X MIC concentrations reduced *Mtb* bacterial burden by 0.3 log_10_ units after 5 days of treatment (P < 0.05; **Fig 1D**). Lack of dose-dependent effects is likely due to either the saturation of the transport machinery available for the drug entry in the macrophages or 1X MIC caused the maximum inhibition possible. The standard TB drug isoniazid reduced the bacterial burden by 1.2 log_10_ units (P < 0.001; **Fig 1D**). To confirm that the decreased bacterial burden in macrophages was due to drug anti-mycobacterial activity and not cytotoxic activity of DON, we evaluated the viability of macrophages in the presence of these same drugs and found no decrease in macrophage viability by MTS assay (**Fig S1**). These results show that DON possesses direct antibacterial activity against *Mtb*.

### 2. JHU083 reduced disease burden in murine TB models

We next evaluated the therapeutic efficacy of JHU083 in a murine model of TB disease. We challenged 129S2 mice with ∼275 CFU of *Mtb* H37Rv via the aerosol route, and randomized mice into three groups: **(1)** untreated, **(2)** JHU083 treatment (1 mg/kg daily), and **(3)** RIF treatment (1.25 mg/kg daily) (**Fig 2A**) with treatment initiation one day after *Mtb* challenge. Lungs were harvested from all three groups at weeks 2 and 5, homogenized, and plated on 7H11 agar plates to quantify the bacillary burden. While JHU083 did not reduce lung colony forming units (CFU) at week 2, it reduced lung bacterial burden by 1.9 log_10_ units after 5 weeks of treatment compared to untreated mice (P < 0.05; **Fig 2B**). To assess JHU083 activity in a mouse model which develops necrotic granulomas, we repeated the experiment in C3HeB/FeJ mice and found that JHU083 treatment for 4 weeks reduced lung bacterial burden by 1.0 log_10_ unit compared to untreated mice (P < 0.05; **Fig S2**). Consistent with lowered bacterial burden, JHU083 treatment significantly prolonged 129S2 mouse survival with a median time to death (MTD) of greater than 60 days compared with untreated controls which showed an MTD of 35.5 days (P <0.01; **Fig 2C**). JHU083-treated mice gained weight to a greater degree than untreated controls during treatment (P <0.0001; **Fig 2D**) and had consistently lower lung weights than the untreated mice (P < 0.001; **Fig 2E**). Quantitative histopathological examination of lungs after 4 weeks of treatment confirmed significant reduction in the granulomatous lesions in that JHU083-treated mice compared to the untreated mice (P < 0.05; **Fig 2F & 2G**). These observations demonstrate that JHU083 inhibits *Mtb* growth in lungs, prolongs survival, and reduces lung pathology in murine TB models.

**Fig 2.**
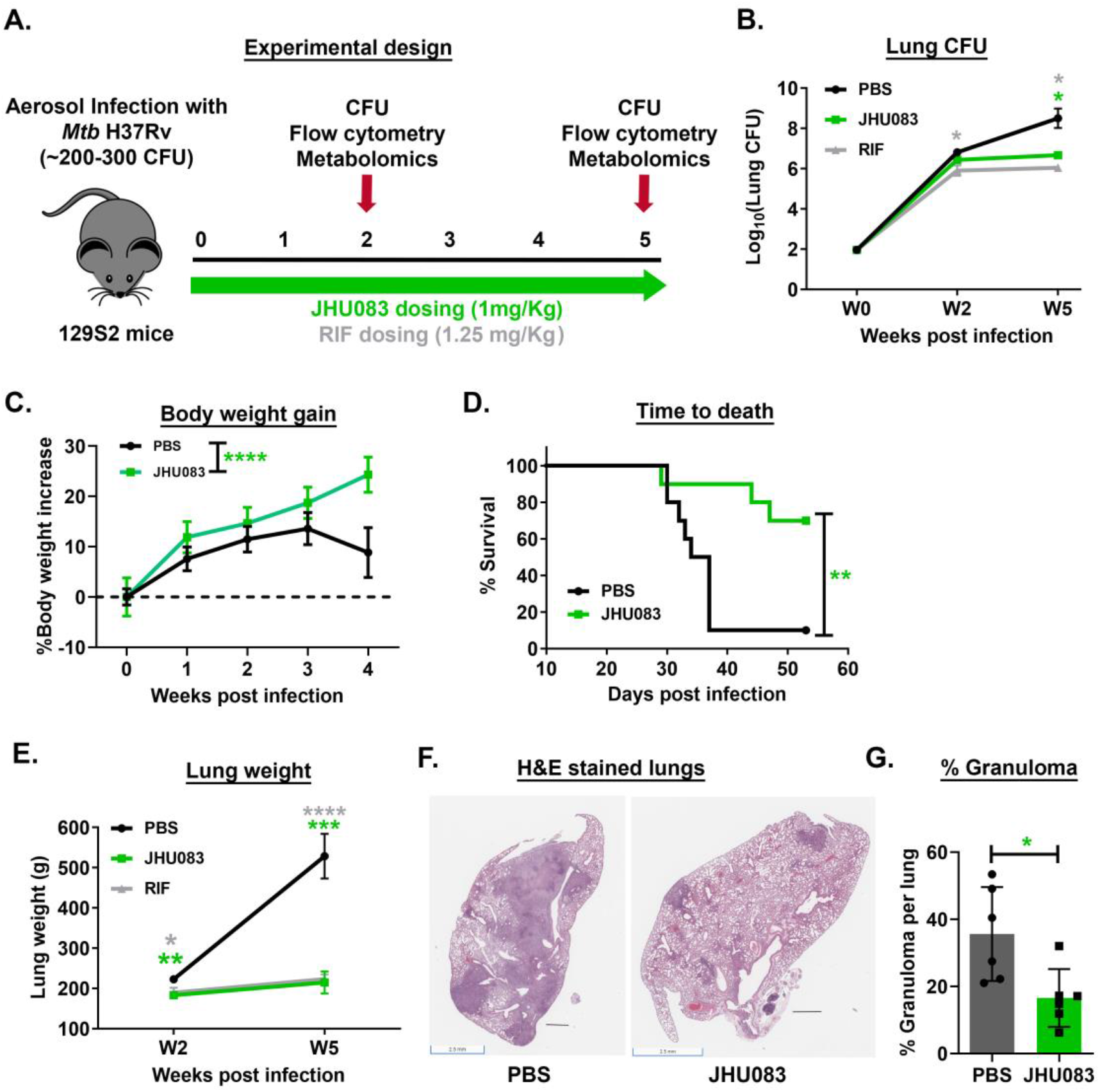
JHU083 administration prevents *Mtb* proliferation in vivo. **(A)** Schematic of the in vivo challenge experiment. 129S2 or C3HeB/FeJ mice (n=4-5) were aerosol infected with ∼200-300 CFU of *Mtb H37Rv*. The mice were treated with JHU083 or RIF via oral route one day after infection. 1 mg/Kg JHU083 was given daily for the first five days, and then the dose was reduced to 0.3 mg/Kg daily (M-F). (**B)** The mice were sacrificed at day 0, weeks 2 and 5 post-infection/treatment. The lungs were harvested, homogenized, serially diluted, and plated on 7H11 selection plates. After 21-25 days, the colonies were counted, and counts were transformed into log_10_ values and plotted. (**C)** The graph shows the percent increase in the body weight of the C3HeB/FeJ mice compared to day 0 with and without JHU083 treatment. (**D)** The survival of control and treated 129S2 mice during the experiment. **(E)** Gross lung weight of 129S2 mice at weeks 2 and 5 post infection/treatment. (**F)** The histopathology of the lungs isolated from C3HeB/FeJ mice infected with *Mtb* H37Rv at week 4 post-infection/treatment. The lungs were formalin fixed, sectioned, and stained with hematoxylin and eosin. (**G)** Quantitation of the lung granuloma areas in the C3HeB/FeJ mice infected with *Mtb* H37Rv at week 4 post-infection/treatment (n=6). Both total granuloma (GA) and lung (LA) areas were measured using ImageScope software (Leica). The percent granuloma (%GA) was calculated using the formula (%GA = (GA X 100)/LA). Data were plotted as Mean ± SEM. Statistical significance was calculated using a two-tailed student t-test considering unequal distribution. For survival study, the drug dosing was discontinued after 6 weeks. For survival curve, log-rank (Mantel-Cox) and Gehan-Breslow-Wilcoxon tests were used and yielded similar p-value. *<0.05, **<0.01, ***<0.001, ****<0.0001. All the experiments were repeated at least two-times.

### 3. JH083 shows no therapeutic efficacy against Mtb in immunocompromised mice

While the Alamar blue assay suggested that JHU083 has direct antibacterial activity against *Mtb*, murine cancer models have shown that JHU083 also exerts beneficial immunomodulatory effects ^6,7^. To delineate its precise mechanism of action, we evaluated the therapeutic efficacy of JHU083 in immunocompromised mice, *Mtb*-infected Balb/c SCID mice. We infected SCID mice with ∼40 CFU of *Mtb* H37Rv on day 0, and individual groups were treated with PBS, JHU083 (1 mg/kg daily), or RIF (1.25 mg/kg daily) starting on day 1(**Fig 3A**). After 5 weeks of infection and treatment, we observed no reduction in the lung bacillary burden in the JHU083-treated mice compared to untreated controls while RIF treatment reduced the lung *Mtb* CFU burden by 4.24 log_10_ units (**Fig 3B**). Additionally, both untreated and JHU083-treated SCID mice showed significant weight loss, while RIF-treated mice continued to gain weight (P < 0.01, **Fig. 3C**). Importantly, JHU083-treated SCID mice died at the same median time to death as untreated mice (MTD 41days), while RIF significantly prolonged survival with an MTD of 73 days (P < 0.0001; **Fig. 3D**). These observations strongly suggest that an intact immune system is required for the therapeutic efficacy of JHU083 against *Mtb*. Despite the drug’s direct antibacterial activity, these data show that JHU083 is likely to exert most of its therapeutic benefit via immunomodulation rather than direct bacterial killing.

**Fig 3.**
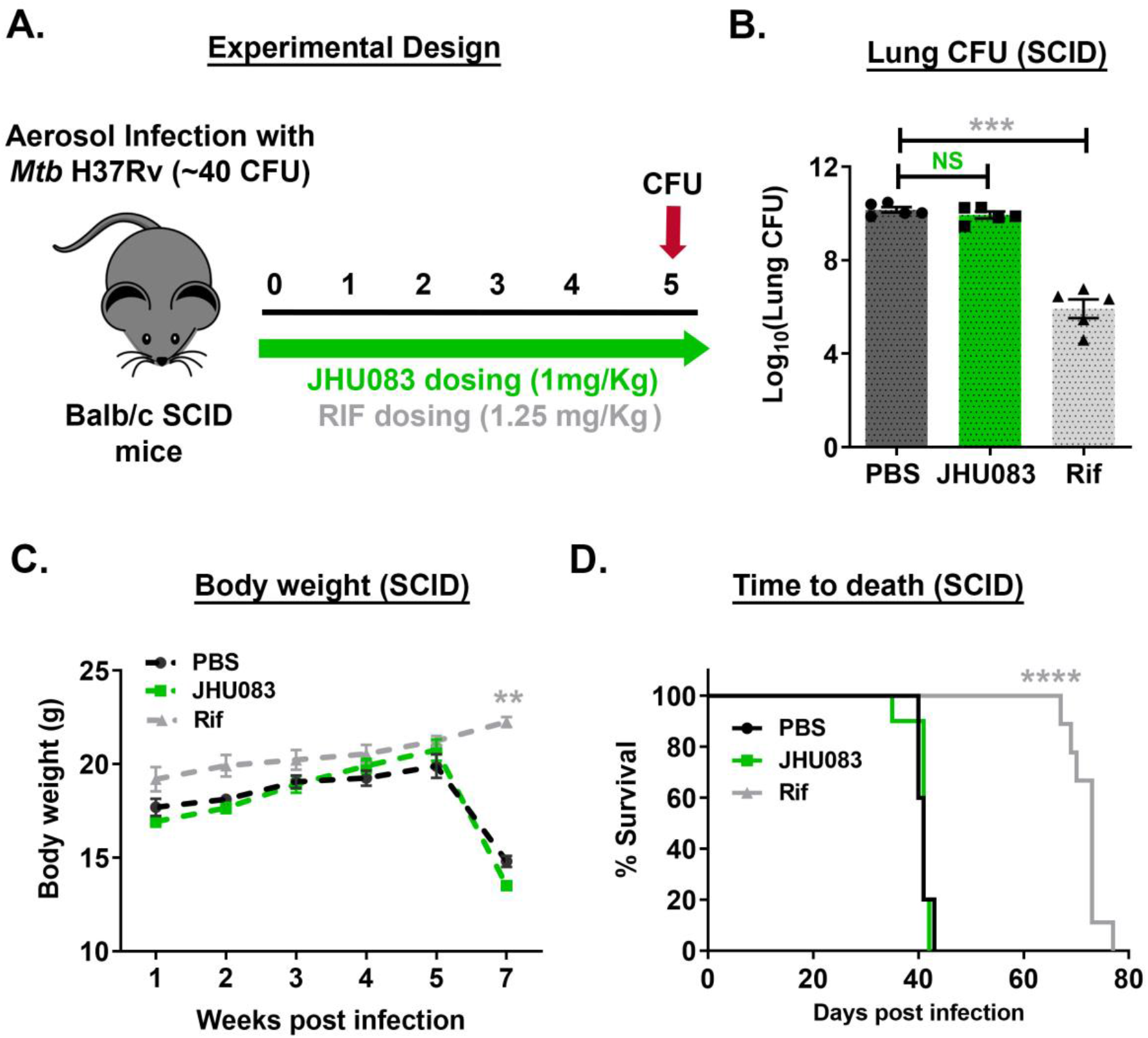
JHU083 loses therapeutic efficacy in *Mtb*-infected immunocompromised mice. **(A)** Schematic of the in vivo experiment. B, Balb/c SCID mice (n=5) were aerosol infected with ∼40 CFU of *Mtb H37Rv*. The mice were then treated with JHU083 or RIF via oral route one day after infection. 1 mg/Kg JHU083 was given daily for the first 5 days and then the dose was reduced to 0.3 mg/Kg daily (M-F) and given for 4 more weeks. (**B)** The mice were sacrificed at week 5 post-infection/treatment. The lungs were harvested, homogenized, serially diluted, and plated on 7H11 selection plates. After 21-25 days, the colonies were counted, and counts were transformed into log_10_ values and plotted. (**C)** The graph shows the effect of both the PBS and Rif controls, and the drug treatment groups on the body weights of SCID mice during the experiment. (**D)** The graph shows the survival of SCID mice during the experiment. Data were plotted as Mean ± SEM. Statistical significance was calculated using a two-tailed student t-test considering unequal distribution. For survival curve, log-rank (Mantel-Cox) and Gehan-Breslow-Wilcoxon tests were used and yielded similar p-value. *<0.05, **<0.01, ***<0.001, ****<0.0001. The experiment was repeated two times.

### 4. Glutamine metabolism inhibition leads to early recruitment of activated T-cells to the lungs during Mtb infection

Several studies in the cancer field have reported that the immunomodulatory effects of glutamine metabolism inhibitors (JHU083 included) are driven by their impact on T-cells ^6,7,20^. Specifically, Leone et al. showed that JHU083 metabolically reprograms T-cells and increases the frequency of long-lived, activated effector T-cells in murine cancer models ^6^. We, therefore, asked if JHU083 would improve effector T-cell immune responses in *Mtb*-infected 129S2 mice. To examine this, we harvested lungs from untreated, JHU083-treated, and RIF-treated mice groups and performed multicolor flow cytometry as described in Methods. Various T and B-cell subsets were identified, as shown in **Fig S3, S4, and S5**. At week 2 post-infection and treatment, the lungs from JHU083-treated mice showed a statistically significant 25% higher frequency of CD4^+^ T-cells (mean = 36.9% of live CD45^+^) compared to both untreated (mean = 29.5% of live CD45^+^) and RIF-treated mice (mean = 24.5% of live CD45^+^) (**Fig 4A**). These CD4^+^ T-cells also showed significant enrichment for the cytokines released by activated T-cells, such as tumor necrosis factor (TNFα) and interferon-gamma (IFNγ) (**Fig 4B and 4C**) and a significant decrease in expression of the immunosuppressive marker interleukin-10 (IL-10; **Fig 4D**). We also found that CD4^+^ T-cell expansion in JHU083-treated mice was driven by significant increases in the frequencies of naïve (CD4^+^ CD62L^+^ CD44-) and follicular helper T-cells (CD4^+^ BCL6^+^) compared to untreated and RIF-treated controls (**Fig 4E & 4F**). Accordingly, both CD4^+^ and CD8^+^ T-cells exhibited an increase in the expression of CD62L and BCL6 in JHU083-treated mice (**Fig 4G-4H & S6A-S6B**). However, there was no overall increase in the frequency of CD8^+^ T-cells (**Fig S6C**). Consistent with naïve T-cell phenotype, both CD4^+^ and CD8^+^ T-cells had lower expression of Klrg1, a terminal differentiation marker, in the JHU083-treated group (**Fig 3I & 3J**). In keeping with an increase in follicular helper T cells, we also found higher frequencies of mature and memory B-cells in both JHU083- and RIF-treated animals compared to untreated controls (**Fig 3K & 3L**). Interestingly, most of the differences in T-cell immune responses in the JHU083-treated group subsided by week 5 (**Fig S7A-S7J)**, indicating that JHU083 had an early but transient effect on T-cell recruitment to *Mtb*-infected lungs.

**Fig 4.**
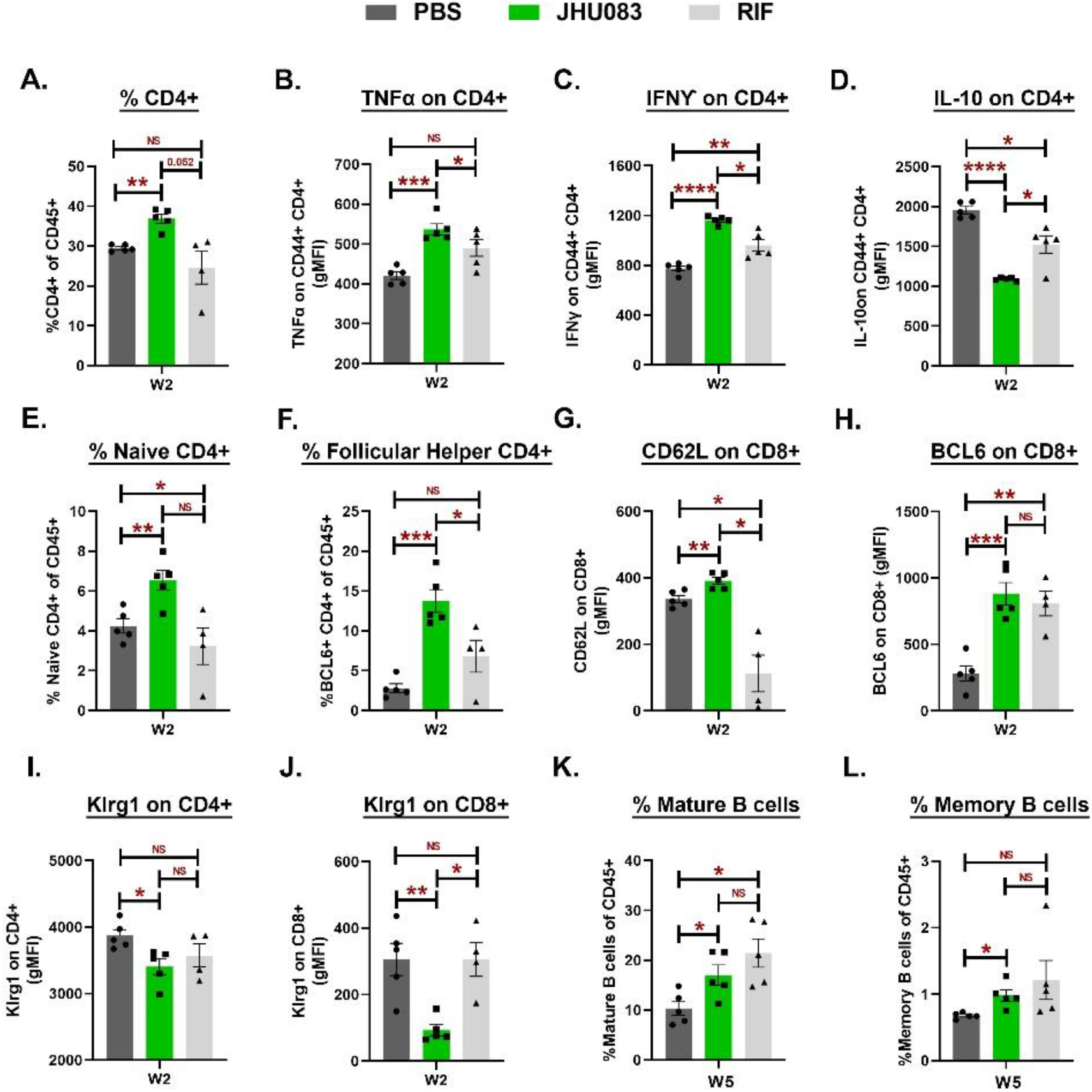
JHU083 administration initiates earlier recruitment of T-cells in the lungs. As described in Fig 2A, *Mtb*-infected 129S2 mice were treated with JHU083 and RIF daily starting day 1 post infection. The mice were sacrificed on weeks 2 and week 5, and the lungs were harvested. Single cell suspensions of the lungs from all three groups were stained with appropriate antibodies and analyzed using multicolor-flow cytometry (n=4-5). We found differences in the **(A)** CD4^+^ T-cell frequency, **(B)** TNFα expression upon activated CD4^+^ T-cells, **(C)** IFNƳ expression upon activated CD4^+^ T-cells, **(D)** IL-10 expression upon activated CD4^+^ T-cells, **(E)** Naïve CD4^+^ T-cells, **(F)** Follicular helper T-cells, **(G)** CD62L expression on CD8^+^ T-cells, **(H)** BCL6 expression on CD8^+^ T-cells, **(I)** Klrg1 expression on CD4^+^ T-cells, **(J)** Klrg1 expression on CD8^+^ T-cells, **(K)** frequency of Mature B-cells, **(L)** frequency of memory B-cells. The X-axis stands for the timepoint at which the lungs were harvested for the flow cytometry experiment. All T-cells data (**A-J**) came from the lungs harvested at week 2 (W2) after infection and treatment while B-cell data (**K-L**) was generated from lungs harvested at week 5 (W5) post-infection and treatment. Data were plotted as Mean ± SEM and are shown as the frequency of CD45^+^ population. gMFI stands for geometric mean fluorescence intensity and was used to define the expression of the individual markers upon the indicated cell types. Statistical significance was calculated using a two-tailed student t-test considering unequal distribution. *<0.05, **<0.01, ***<0.001, ****<0.0001. The experiment was repeated two times.

### 5. Glutamine metabolism inhibition modulates macrophages towards an inflammatory signature

Myeloid cells, including macrophages and myeloid-derived suppressor cells (MDSCs), are known to play an important role in *Mtb* containment ^14^. Since JHU083 is also known to reprogram myeloid cells and reinforce their host-protective functions in murine cancer models, we investigated the impact of JHU083 administration upon lung myeloid cell populations in the mouse TB model. Using the same model in 129S2 mice (**Fig 2A**), we performed flow cytometry on lung cells from mice sacrificed at weeks 2 and 5 post-infection/treatment as described in Methods. The gating strategy is described in **Fig S8 and S9**.

We found that total lung CD11b^+^ myeloid cells were significantly increased at week 5, but not at week 2 in both JHU083- and RIF-treated mice compared to untreated mice (**Fig 5A**). We then evaluated three major myeloid cell subsets: **(1)** alveolar macrophages (AM; CD45^+^ SiglecF^+^); **(2)** Interstitial macrophages (IM; CD45^+^ CD11b^+^ F4/80^+^), and **(3)** myeloid-derived suppressor cells (MDSCs; CD45^+^ CD11b^+^ Ly6G/Ly6C). We detected a two-fold decrease in the frequency of AM in JHU083-treated mice compared to untreated mice at week 2, but not at week 5 (**Fig 5B**, P = 0.05). At the 2-week timepoint, the JHU083-treated group exhibited higher expression of the co-stimulatory molecule CD86 (**Fig 5C**) and lower expression of inhibitory receptor CD206 than untreated controls (**Fig 5D**), consistent with the AMs present in the lung at 2 weeks being enriched for a proinflammatory phenotype.

**Fig 5.**
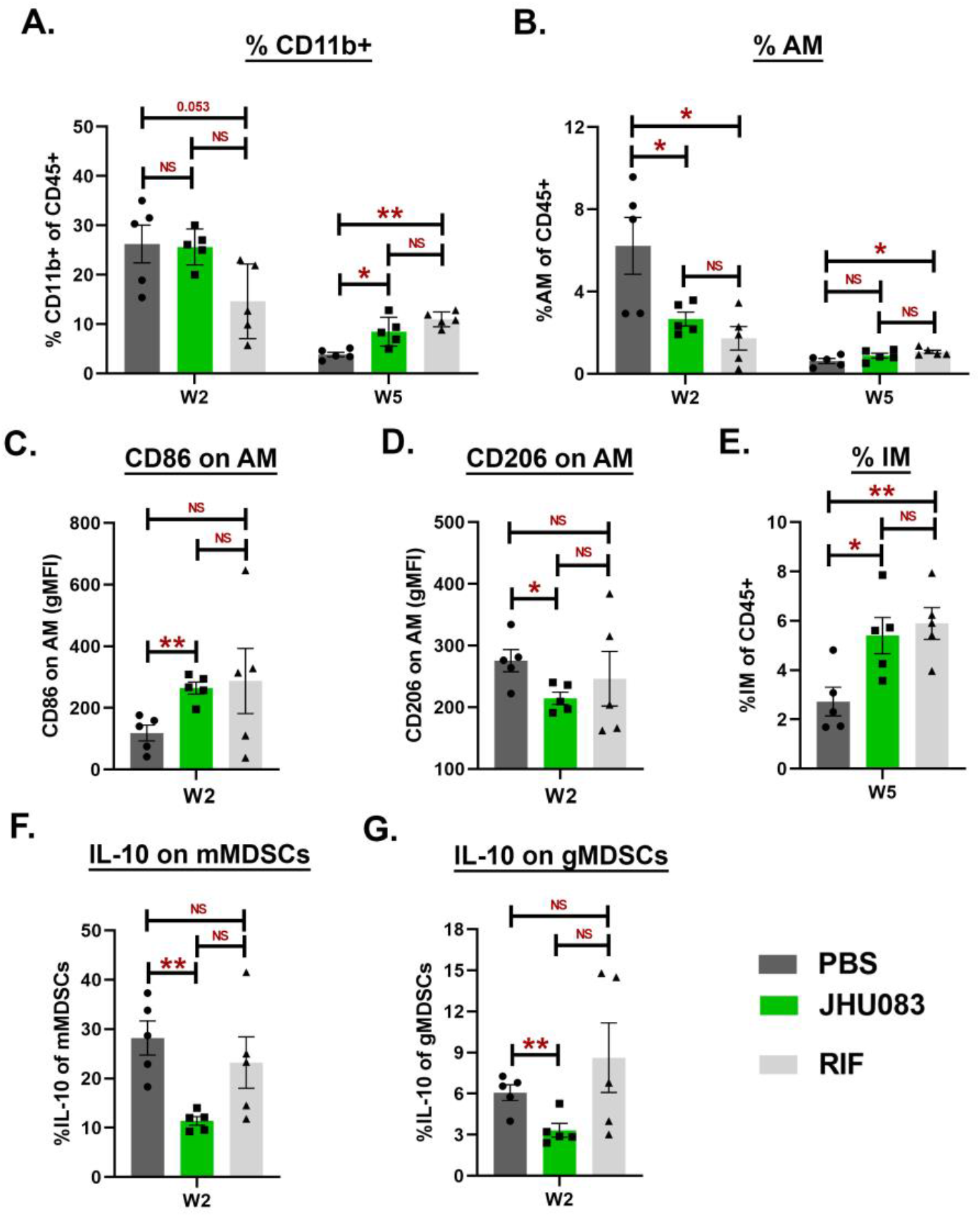
JHU083 treatment causes infiltration of the proinflammatory myeloid cells in the lungs. As described in Fig 2A, *Mtb*-infected 129S2 mice were treated with JHU083 and RIF every day starting day 1 post-infection. The mice were sacrificed on weeks 2 and week 5, and the lungs were harvested. Single cell suspension of the lungs from all three groups were stained with appropriate antibodies and analyzed using multicolor-flow cytometry (n=5). Details are provided in “Methods” section. We found differences in the **(A)** CD11b^+^ myeloid cells, **(B)** alveolar macrophages (AM), **(C)** CD86 expression upon AM, **(D)** CD206 expression upon AM, **(E)** Interstitial macrophages (IM), **(F)** IL-10 expression upon monocytic MDSCs (mMDSCs), **(G)** IL-10 expression upon granulocytic MDSCs (gMDSCs). The X-axis shows the timepoint at which the lungs were harvested for flow cytometry experiment. Data were plotted as Mean ± SEM and are shown as the frequency of CD45^+^ population. gMFI stands for geometric mean fluorescence intensity and was used to define the expression of the individual markers upon the indicated cell types. Statistical significance was calculated using a two-tailed student t-test considering unequal distribution. *<0.05, **<0.01. The experiment was repeated two times.

In contrast to AMs, the IM cell frequencies were unchanged in JHU083-treated animal at 2 weeks compared to uninfected animals, but significantly increased (2-fold) in JHU083-treated animals at the late timepoint of week 5 (**Fig. 5E, Fig S10A**). In spite of this late recruitment of IMs to the lungs in JHU083-treated mice, the expression of either CD86 or CD206 remained constant, suggesting a balance of M1- and M2-like IMs entering the lungs (**Fig. S10B and S10C)**.

We then analyzed two prominent subsets of myeloid-derived suppressor cells (MDSCs) in these murine lung samples: **(1)** Monocytic MDSCs (mMDSCs; CD11b^+^ Ly6G-Ly6C^High^) and **(2)** Granulocytic MDSCs (gMDSCs; CD11b^+^ Ly6G^+^ Ly6C^Low^). mMDSCs were present at significantly higher frequencies at both weeks 2 and 5 compared to the untreated group (**Fig. S10D**), while gMDSCs did not significantly differ in the JHU083-treated animal compared with the other groups (**Fig. S10E**). However, we found that both IL-10-expressing mMDSCs and gMDSCs were present in reduced frequencies at the early 2-week timepoint, but subsequently were not significantly different from the other groups at 5 weeks (**Fig. 5F, 5G, S10F and S10G**). Thus, our analysis of lung myeloid cells revealed an early reduction of *Mtb*-permissive AMs which later normalized, a late influx (5 weeks) of *Mtb*-restrictive IMs, and an early reduction of immunosuppressive subtypes, namely IL10^+^ mMDSCs, and IL-10^+^ gMDSCs. These myeloid cell shifts mediated by JHU083-treatment suggest that the drug promotes enrichment of cells restrictive of *Mtb* growth and reduces those associated with immunosuppression.

### 6. JHU083 drives host-protective metabolic changes in Mtb-infected lungs

Since JHU083 has been shown to reprogram metabolic pathways of both cancer and immune cells by blocking glutamine metabolism^6,7^, we hypothesized that JHU083 administration would alter metabolic pathways in *Mtb*-infected lungs. To investigate this, we performed LC/MS-based metabolomics of total lung tissues from untreated, JHU083- and RIF-treated animals at weeks 2 and 5. We detected a total of 144 metabolites and found alterations in the levels of >100 metabolites among the groups pointing toward a complex metabolic reprogramming (**Supplementary Table S1**). JHU083-treated mice exhibited maximal changes in metabolite levels at week 2 post-infection and treatment (**Fig. S11**) in contrast to the RIF-treated group that showed minimal changes at week 2 (**Fig. S12)**. The most notable changes in JHU083-treated lungs were as follows: (**i)** A **∼**17% reduction in the glutamine levels (P = 0.17; **Fig. 6A**), **(ii)** Alterations in the metabolism of arginine with accumulation of citrulline (1.4-fold increase; P = 0.04; **Fig. 6B**) suggesting greater activity of nitric oxide synthase (NOS) and the likely generation of the host-protective molecule NO. This was accompanied by significantly lower levels of other arginine metabolites including ornithine, polyamines, S-adenosyl-homocysteine, creatine, and agmatine (**Fig. 6E, S13A**). **(iii)** Alterations in the metabolism of tryptophan with accumulation of 5-hydroxy-3-indoleacetic acid (5-HIAA; 2.9-fold increase, P = 0.008; **Fig. 6C**) and ∼60% reduction in the level of quinolinic acid which is a byproduct of the immunosuppressive metabolite kynurenine (P = 0.02; **Fig. 6D, 6F, S13B**). These data show that JHU083-treatment leads to lower glutamine levels in the lung though the change was modest and not statistically significant. Importantly, however, JHU083 treatment altered the metabolism of two immunologically important amino acids, arginine, and tryptophan, in directions that may drive enhanced anti-*Mtb* host immune responses (**Fig. 7**).

**Fig 6.**
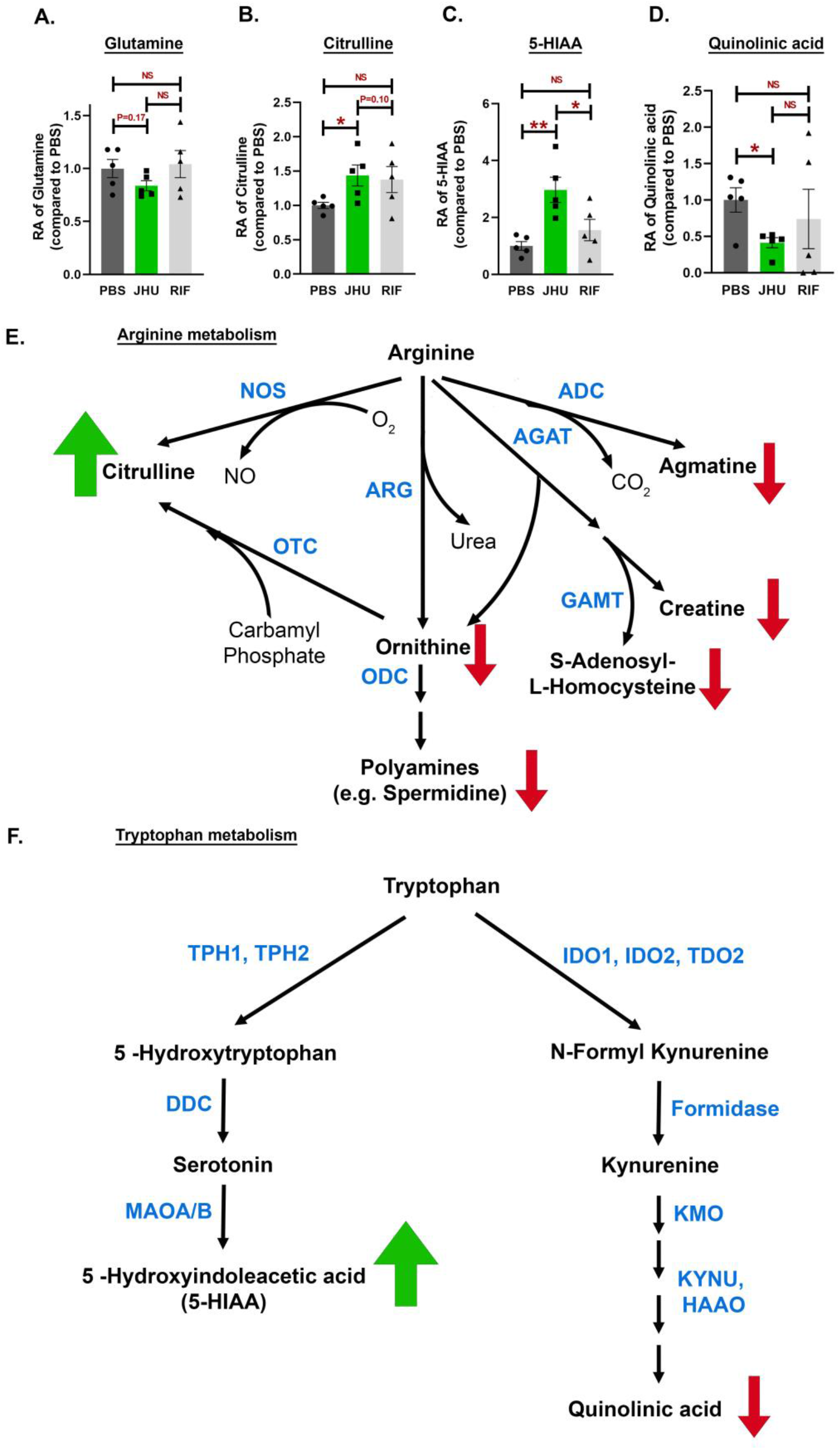
JHU083 treatment drives metabolic reprogramming in the *Mtb*-infected lungs. As described in Fig 2A, *Mtb*-infected 129S2 mice were treated with JHU083 and RIF every day starting day 1 post-infection. Mice were sacrificed at week 2, the lungs were harvested, and total metabolites were methanol extracted as described in “Methods”. The total metabolites were normalized to the tissue weight and then to the untreated controls. We detected changes in the level of **(A)** glutamine, **(B)** Citrulline, **(C)** 5-HydroxyIndole acetic acid (5-HIAA) and (**D)** Quinolinic acid. Schematic representation of the **(E)** arginine and **(E)** tryptophan metabolism listing the metabolic steps and the metabolites relevant to the study. Green and red arrows next to a metabolite indicate accumulation and depletion respectively in JHU083-treated group compared to untreated control. Abbreviations stands for; Arginine decarboxylase (ADC); L-arginine:glycine amidinotransferase (AGAT); Arginase (ARG); Guanidinoacetate N-methyltransferase (GAMT); Nitric oxide synthase (NOS); Ornithine decarboxylase (ODC); Ornithine decarboxylase (OTC); Dopa decarboxylase (DDC); 3-hydroxyanthranilate 3,4-dioxygenase (HAAO); Indoleamine 2,3-dioxygenase (IDO); Kynurenine 3Monooxygenase (KMO); Kynureninase (KYNU); Monoamine oxidase A/B (MAOA/B); Tryptophan 2,3dioxygenase (TDO2); Tryptophan hydroxylase (TPH1/2). Data were plotted as Mean ± SEM. Statistical significance was calculated using a two-tailed student t-test considering unequal distribution. *<0.05, **<0.01. The experiment was repeated two times.

**Fig 7.**
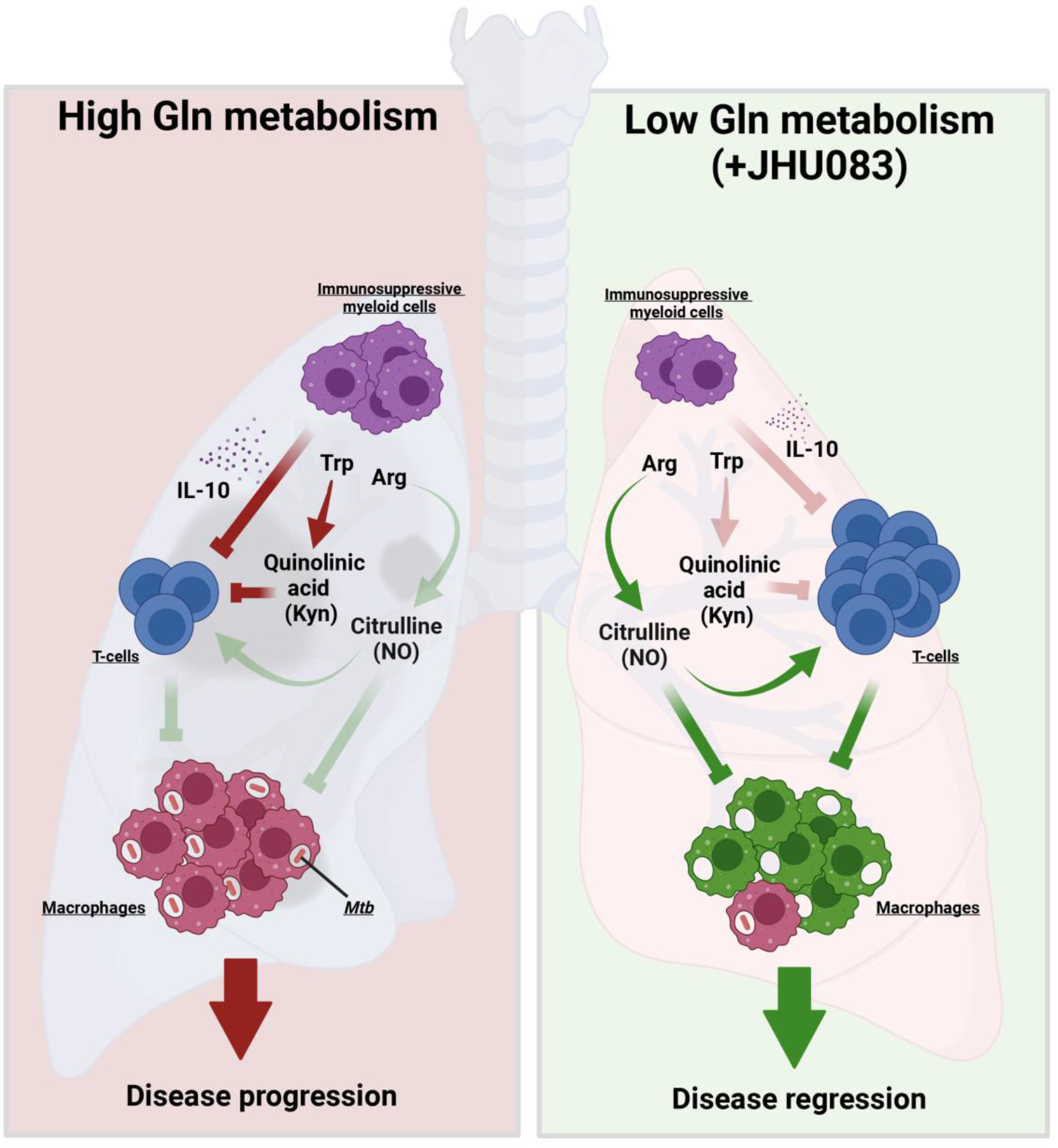
Model depicting the mechanism of action of JHU083. Under glutamine-sufficient conditions, IL-10 produced by immunosuppressive myeloid cells (MDSCs) and quinolinic acid (a product of the Kynurenine pathway) inhibit T-cell proliferation ad functions, promoting infection and disease progression. Glutamine antagonist JHU083, decreases IL-10-producing MDSCs leading to a higher frequency of T-cells. Depletion of quinolinic acid and accumulation of citrulline further potentiates T-cell functions while preventing *Mtb* proliferation in macrophages. These immunometabolic changes then lead to disease regression and improved lung histology. Red lines represent host-deleterious process, while green lines are for host-protective processes.

## DISCUSSION

*Mtb*, a sophisticated intracellular pathogen, deploys numerous strategies to overcome the host immune system, starting from interfering with immune cell recruitment, inhibiting their host-protective functions and compromising their metabolic fitness. Interestingly, little is known about the effect of *Mtb* infection upon host metabolic pathways or how these alterations contribute to TB pathogenesis. Here, we show that JHU083-mediated glutamine metabolism inhibition prevents *Mtb* proliferation both in vitro and in vivo. JHU083-treated *Mtb*-infected mice had lowered lung bacillary burden, gained weight, and lived longer with improved lung histology. JHU083 treatment also caused an earlier onset of T-cell recruitment and a reduced frequency of immunosuppressive myeloid cells in the lungs. Metabolomics analysis revealed that JHU083-treated lungs accumulated citrulline (suggesting greater host-protective NOS activity) and lowered quinolinic acid (a byproduct of the immunosuppressive molecule kynurenine). Overall, JHU083-treated animals exhibited an improved anti-*Mtb* immune response.

JHU083 is a prodrug that is converted to 6-diazo-5-oxo-norleucine (DON) in the serum and the tissues in the presence of esterases and peptidases. DON exhibits high structural similarity to glutamine, blocks several reactions that either generate or require glutamine and has been extensively studied as a potential cancer therapeutic^12,21^. However, initial clinical trials were hampered by the dose-limiting gastrointestinal (GI) toxicity of DON ^12^. The prodrug strategy has been shown to not only reduce GI toxicity but also increase bioavailability ^8,11,13^. In the tumor microenvironment, elevated glutamine metabolism has been associated with increased immunosuppression leading to a blunted effector T-cell immunity and the promotion of tumor growth ^9,22,23^. JHU083-mediated glutamine antagonism reduces immunosuppression in the tumor microenvironment by enhancing the recruitment of long-lived proliferating T-cells, thereby inhibiting tumor growth ^6,7^. The antitumor effects of JHU083 have been demonstrated in multiple tumor models including glioma, colon, lung, and triple-negative breast cancers ^6,7,11,24^. A prodrug closely related to JHU083, DRP104 (Sirpiglenastat) is in early-phase clinical trials for solid tumors (ClinicalTrials.gov Identifier: NCT04471415) ^11^. In addition, DRP104 already has received US FDA Fast Track designation for the treatment of non-small cell lung cancer patients.

In contrast to cancer, the literature on the importance of host glutamine metabolism in infectious diseases such as TB is limited. Specifically, there is no consensus on the impact of exogenous glutamine levels to reduce the progression of a constellation of respiratory diseases ^25^. In the field of acute lung injury, a few studies have shown a beneficial effect of Gln supplementation ^26,27^, while others have deemed Gln inhibition to be important for preventing progression following injury ^28^. In the case of TB, *Mtb* infection has been shown to induce glutamine metabolism transcripts ^29,30^. Several studies have found that glutamine metabolism is crucial for T-cell cytokine production, M1 polarization of macrophages, and that glutamine serves as an important carbon and nitrogen source in *Mtb-*infected macrophages ^29,31,32^. Thus, while previous reports have described potential roles for host glutamine metabolism during TB, direct inhibition of host glutamine metabolism as a potential host-directed therapy for TB has not been explored previously.

Interestingly, *Mtb* glutamine synthetases have been extensively studied as antibacterial drug targets. Glutamine synthetase catalyzes the amidation of glutamate to glutamine in an ATP-dependent reaction ^17^. *Mtb* possess four such enzymes, yet only GlnA1 has been shown to be secreted and to be essential for in vitro and in vivo growth of *Mtb* ^16^. In one study, a GlnA1 peptide was found in the serum of 82% latent TB patients ^33^. GlnA1 inhibition using methionine sulfoximine (MSO), an inhibitor of glutamine synthases, lowered lung bacillary burden in *Mtb*-infected guinea pigs ^15,34^. However, the initial enthusiasm regarding GlnA1 as a potential therapeutic target for *Mtb* infection was dampened by the rapid emergence of spontaneous mutants resistant to MSO (MSO^R^). The mutations were found to be in the upstream promoter region of *glnA1* causing GlnA1 overexpression potentially conferring drug resistance ^19^. In our own hands using MSO-containing agar, we readily isolated spontaneous MSO^R^ mutants. Interestingly, these MSO^R^ mutants showed no cross-resistance to JHU083 indicating differences in their mode of action and cellular targets (unpublished data).

As high glutamine metabolism promotes immune tolerance and immunosuppression in several disease models ^7,35^, we hypothesized that *Mtb*-associated glutamine release is an immunometabolic virulence mechanism that may be countered by a pharmacologic inhibition of local glutamine metabolism. Data presented here support the hypothesis that the inhibition of Gln metabolism has immunomodulatory effects as shown by enhanced levels of effector T-cells, decreased levels of immunosuppressive myeloid cells, and the absence of a JHU083 protective effect in SCID mice. We also performed whole lung metabolomics in the presence and absence of JHU083 and observed that JHU083-treatment elicits metabolic shifts consistent with enhanced anti-*Mtb* host immune responses.

CD4+ T-cells are critical in driving anti-*Mtb* immune responses. T-cells rely on several amino acids (arginine, tryptophan, glutamine etc.) for their proliferation, activation, and optimal function^36^. Immunosuppressive cells (such as MDSCs and Tregs) create an artificial local depletion of these amino acids, which adversely affects T-cell proliferation and function ^14,37^. For example, immunosuppressive myeloid cells overexpress arginase, scavenging arginine and converting it to ornithine and urea ^38^. We found that JHU083 treatment lowered ornithine levels while increasing the level of citrulline in the lung (**Fig 6B**). Citrulline originates from arginine in the presence of inducible nitric oxide synthase (iNOS) and generates nitric oxide (NO). These results suggest that JHU083 treatment potentially produces NO that has host-protective functions. NO has potent anti-mycobacterial activity controlling *Mtb* proliferation in macrophages ^39,40^. Citrulline also serves as an arginine reservoir and contributes to maintaining optimal NO concentration when free arginine is scarce ^41,42^. In addition, citrulline is also known to augment T-cell proliferation and functions ^43^. Diminished levels of other arginine metabolites such as spermidine, creatine, agmatine are also suggestive of a preferential upregulation of the NOS pathway (**Fig S13A**).

Another crucial amino acid for T-cell division especially during activation is tryptophan^36^. Suppressive myeloid cells starve T-cells of tryptophan by overexpressing IDO1 and by converting tryptophan to kynurenine ^35^. Upregulated IDO expression in tuberculous granulomas has been shown to promote *Mtb* survival in the host, and IDO1 inhibition improves disease outcome ^44-46^. Kynurenine, the product of IDO activity, has been shown to accumulate in the plasma of TB patients ^47,48^. Kynurenine metabolism also generates quinolinic acid, a potent neurotoxin (if accumulated), which is essential for the de novo synthesis of NAD^+ 49^. JHU083-treatment lowered quinolinic acid levels in the lungs without decreasing cellular NAD^+^ levels (**Supplementary Table 1)**. The lowered quinolinic acid levels may be due to either downregulation of tryptophan-kynurenine pathway or heightened rate of quinolinic acid utilization. Interestingly, JHU083 treatment led to accumulation of 5-HIAA, another tryptophan metabolite. 5-HIAA is primary breakdown product of serotonin and is often used as a proxy for the serotonin measurement. Peripheral serotonin that accounts for >90% of the pool has been shown to exert immunomodulatory effects ^50^. Studies have shown that stimulation or activation of serotonin receptor A promotes T-cell proliferation and phagocytic abilities of macrophages ^50^. We are not aware of any study reporting the correlation of 5-HIAA with TB.

There are several similarities in the immunomodulatory activity of JHU083 in both the TB and cancer models. JHU083 treatment increased the frequency of CD4+ T-cells and enhanced naive T-cell signatures in both models ^6,7^. We also found an increased frequency of follicular helper T-cells and mature B-cells. In both TB and cancer, JHU083 administration lowered the abundance of immunosuppressive myeloid cells while increasing the abundance of myeloid cells with proinflammatory signatures. In both models, the lungs of JHU083-treated mice showed reduce tryptophan metabolism towards kynurenine pathway. However, there were a few key differences between the TB and cancer data. First, we did not observe a decline in the overall frequency of MDSCs in JHU083-treated *Mtb*-infected mice. The reduction that we observed was limited to the IL-10-producing MDSCs population. Second, the whole lungs of JHU083-treated *Mtb*-infected mice showed a trend toward diminished glutamine pools while the cancer data showed an accumulation of glutamine in the tumor microenvironment as measured in excised syngeneic, solid tumors taken from the flank. Indeed, in future studies, it will be interesting to evaluate solid granulomatous lesions themselves, rather than the whole lung, to better understand the effects of JHU083 on the granuloma microenvironment.

While we have demonstrated that JHU083 treatment results in both antibacterial and immunomodulatory activities, we propose that the host-directed activity results in the predominant therapeutic effect. In support of this hypothesis, we have shown that **(1)** JHU083 loses its therapeutic efficacy in *Mtb*-infected immunocompromised mice (**Fig 3**), that **(2)** there are significant beneficial changes in lymphoid and myeloid cell populations with JHU083, and that **(3)** these immune cell changes mostly occur at week 2 post-infection and treatment when there is no difference in the lung bacillary burden between the JHU083- and PBS-treated groups (**Fig 2B**). To further delineate antibacterial vs immunomodulatory activities of JHU083, evaluating efficacy of JHU083 in mice infected with an *Mtb* mutant resistant to JHU083 (JHU^R^) would be valuable, and the lack of such data are a limitation of this study. Indeed, we made numerous unsuccessful attempts to isolate resistant mutants on JHU083-containing agar plates although MSO^R^ mutants were readily found by the same method. Another limitation is that the metabolomic data presented in **Fig 6** and **S11-S13** are the result of an unbiased, untargeted (whole lung) analysis of ∼150 cellular metabolites from the lungs of Mtb-infected, JHU083-treated, and untreated mice. Future studies to determine the metabolic status of specific immune cell populations and to measure metabolite fluxes with and without drug treatment will be valuable in refining our current observations.

Based on both the results presented in the prior literature by Horwitz and colleagues ^15-17^ and in this study, we propose that *Mtb* lung infection generates local microenvironments with elevated glutamine levels. This in turn leads to accumulation of immunosuppressive myeloid cells, reduced effector T-cell function, and downregulation of NO and citrulline synthesis. We have shown that administration of JHU083 in the TB mouse model results in the decreased level of immunosuppressive myeloid cells, enhanced levels of effector T-cells, and increased levels of citrulline (and possibly NO) production (**Fig 7**). Lastly, while JHU083 has both a direct antibacterial effect and immunomodulatory activity, the kinetics of its effects and its lack of efficacy in SCID mice suggest that JHU083 functions predominately as a host-directed immunotherapeutic in the mouse model of TB.

## AUTHOR CONTRIBUTIONS

SP, JRM, and WRB conceptualized the study and designed the research approach; BSS, RR, RDL, JDP provided inputs critical to the study; BSS and RR provided JHU083 for the study; SP, JS, SL, BK performed animal studies; BK performed BMDM experiment; SP, JS and LZ performed metabolomics study; SP performed flow cytometry; SP, JRM and WRB analyzed data; SP, JRM and WRB wrote the paper; SP, JS, SL, LZ, BK, RDL, RR, JDP, JRM, BSS and WRB critically reviewed the manuscript.

## ACKNOWLEDGEMENTS

We gratefully acknowledge the support of NIH grant AI155602, R01CA226765 and the Bloomberg-Kimmel Institute for Cancer Immunotherapy. Flow cytometry was performed at the SKCCC flow/mass cytometry core. Lung histology was performed at SKCCC oncology tissue services core. We are grateful to Drs. Marcus Horwitz and Michael Tullius for graciously sharing GlnA1 auxotrophic mutant strains. We acknowledge past and present members of the Bishai lab for helpful suggestions and discussion throughout the study.

---

**Fig S1.**
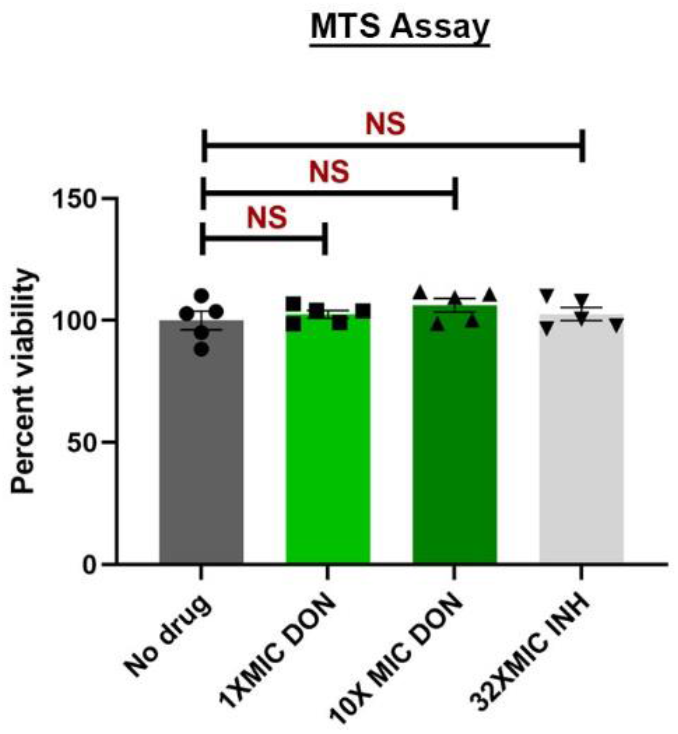
The effect of JHU083 upon the viability of bone-marrow derived macrophages (BDDMs) in the MTS assay. BMDMs were harvested from the femurs of 8-12 weeks old C57BL6 mice, activated using IFNƳ, and then treated with DON and INH. MTS assay was performed after 5 days of daily drug treatment. The experiment was performed in triplicate.

**Fig S2.**
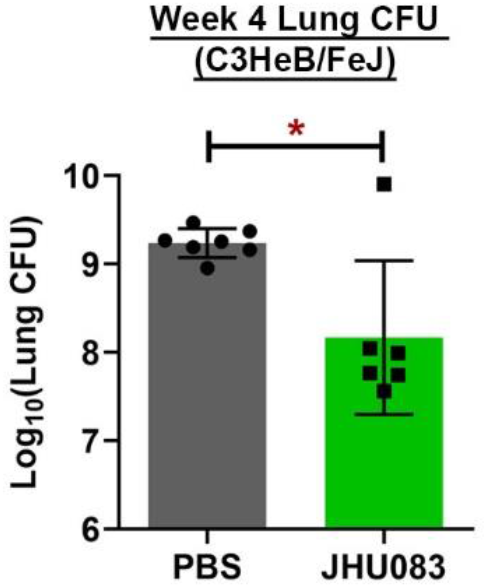
JHU083 administration reduces lung bacillary burden in *Mtb*-infected C3HeB/FeJ mice. C3HeB/FeJ mice (n=4-5) were aerosol infected with ∼150 CFU of *Mtb H37Rv*. The mice were then treated with JHU083 via oral route one day after infection. 1 mg/Kg JHU083 was given daily for the first 5 days and then the dose was reduced to 0.3 mg/Kg daily (M-F). (**B)** The mice were sacrificed at day 0 and week 5 post-infection/treatment. The lungs were harvested, homogenized, serially diluted, and plated on 7H11 selection plates. After 21-25 days, colonies were counted, and counts were transformed into log_10_ values and plotted. Data is plotted as Mean ± SEM. Statistical significance was calculated using a two-tailed student t-test considering unequal distribution. *<0.05. The experiment was performed two-times.

**Fig S3.**
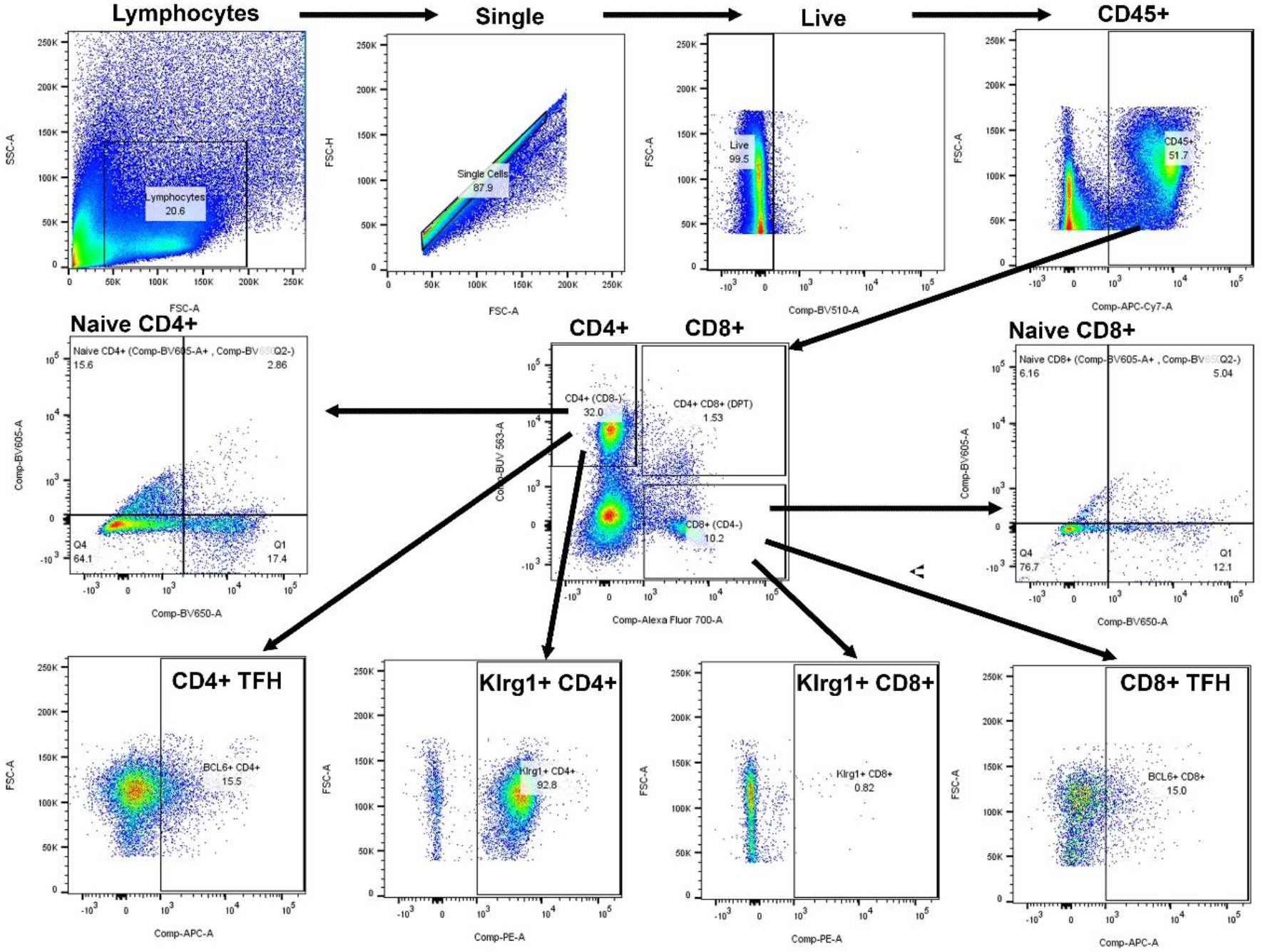
Two-dimensional gating strategies for flow cytometrical identification of T-cell subsets. We excluded doublets and debris and gated on single live CD45^+^ cells. We identified CD4^+^ T-cells (CD45^+^ CD8^-^ CD4^+^), CD8^+^ T-cells (CD45+ CD4^-^ CD8^+^), Naïve CD4^+^ T-cells (CD45^+^ CD8^-^ CD4^+^ CD44^-^ CD26L^+^), Naïve CD8^+^ T-cells (CD45^+^ CD8^+^ CD4^-^ CD44^-^ CD26L^+^), Follicular helper CD4^+^ T-cells (CD45^+^ CD8^-^ CD4^+^ BCL6^+^), Follicular helper CD8^+^ T-cells (CD45^+^ CD8^+^ CD4^-^ BCL6^+^), Klrg1^+^ CD4^+^ T-cells (CD45^+^ CD8^-^ CD4^+^ Klrg1^+^), Klrg1^+^ CD8^+^ T-cells (CD45^+^ CD8^+^ CD4^-^ Klrg1^+^).

**Fig S4.**
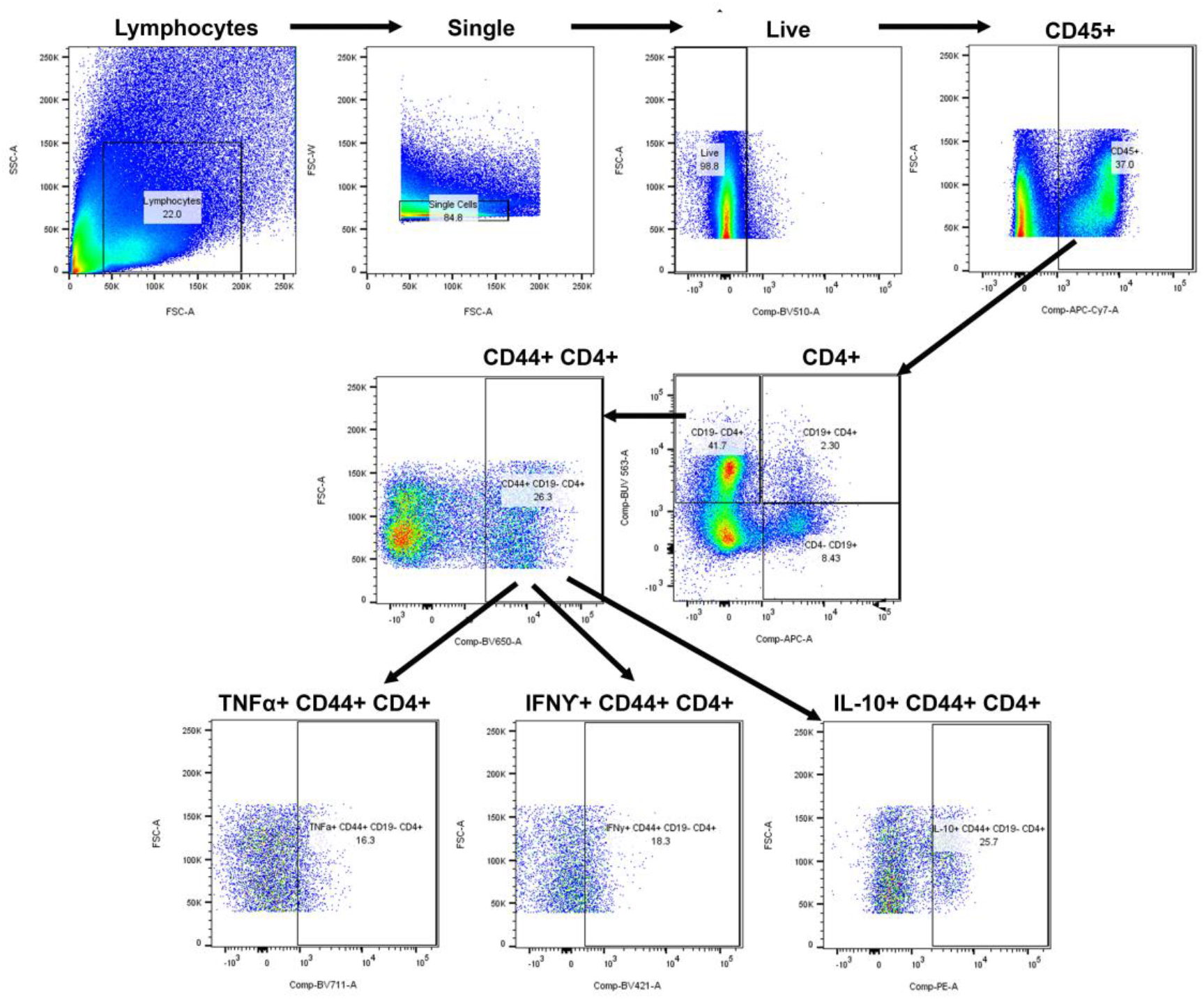
Two-dimensional gating strategies for flow cytometrical identification of cytokine-producing T-cell subsets. We excluded doublets and debris and gated on single live CD45+ cells. We identified CD4^+^ T-cells (CD45^+^ CD19^-^ CD4^+^), activated CD4^+^ T-cells (CD45^+^ CD19^-^ CD44^+^ CD4^+^), TNFα^+^ activated CD4^+^ T-cells (CD45^+^ CD19^-^ CD44^+^ CD4^+^ TNFα^+^), IFNƳ^+^ activated CD4^+^ T-cells (CD45^+^ CD19^-^ CD44^+^ CD4^+^ IFNƳ^+^) and IL-10^+^ activated CD4^+^ T-cells (CD45^+^ CD19^-^ CD44^+^ CD4^+^ IL10^+^).

**Fig S5.**
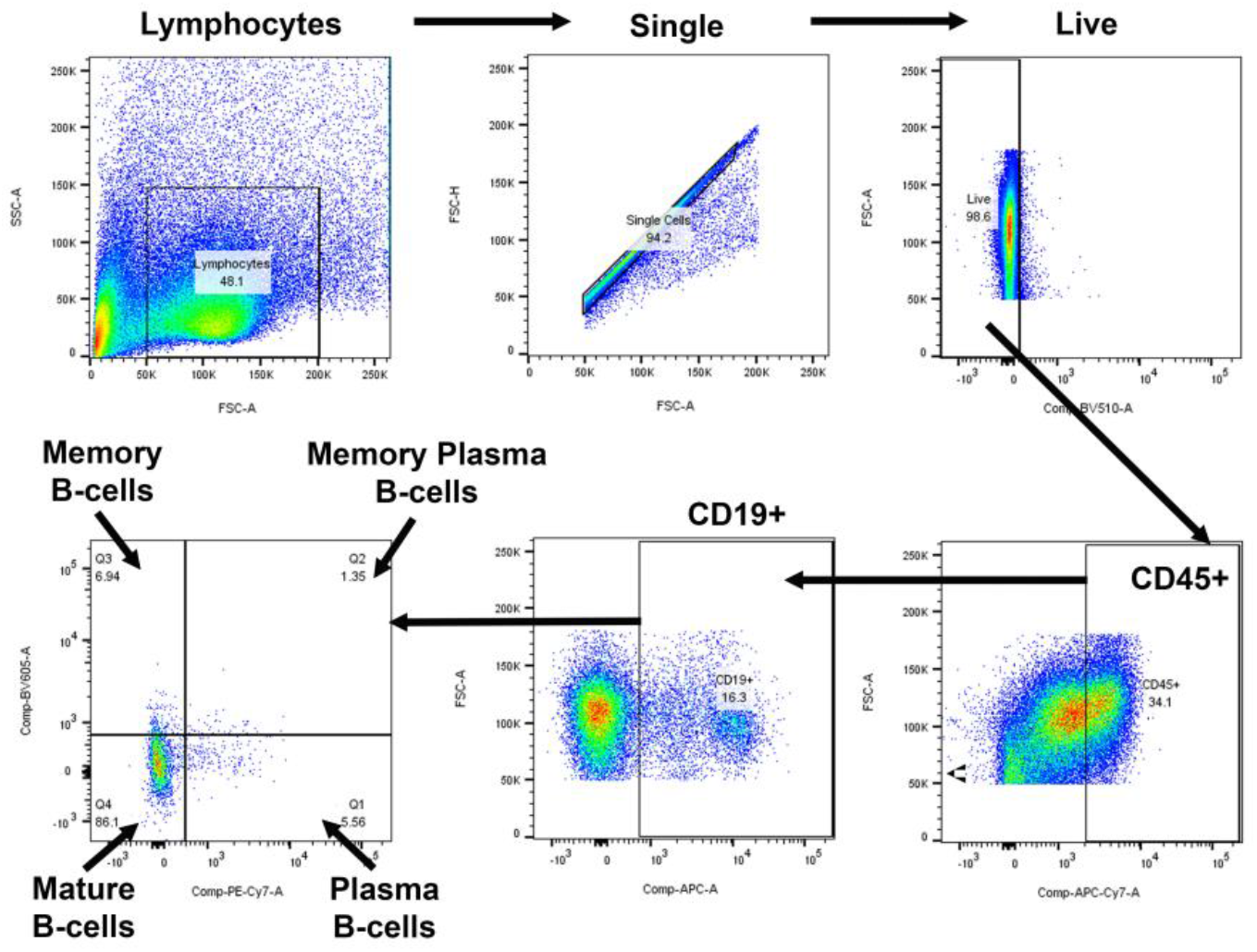
Two-dimensional gating strategies for flow cytometrical identification of B cell subsets. We excluded doublets and debris and gated on single live CD45^+^ cells. We identified total T-cells (CD45^+^ CD19^+^), mature B-cells (CD45^+^ CD19^+^ CD27^-^ CD138^-^), memory B-cells (CD45^+^ CD19^+^ CD27^+^ CD138^-^), plasma cells (CD45^+^ CD19^+^ CD27^-^ CD138^+^), plasma memory B-cells (CD45^+^ CD19^+^ CD27^+^ CD138^+^).

**Fig S6.**
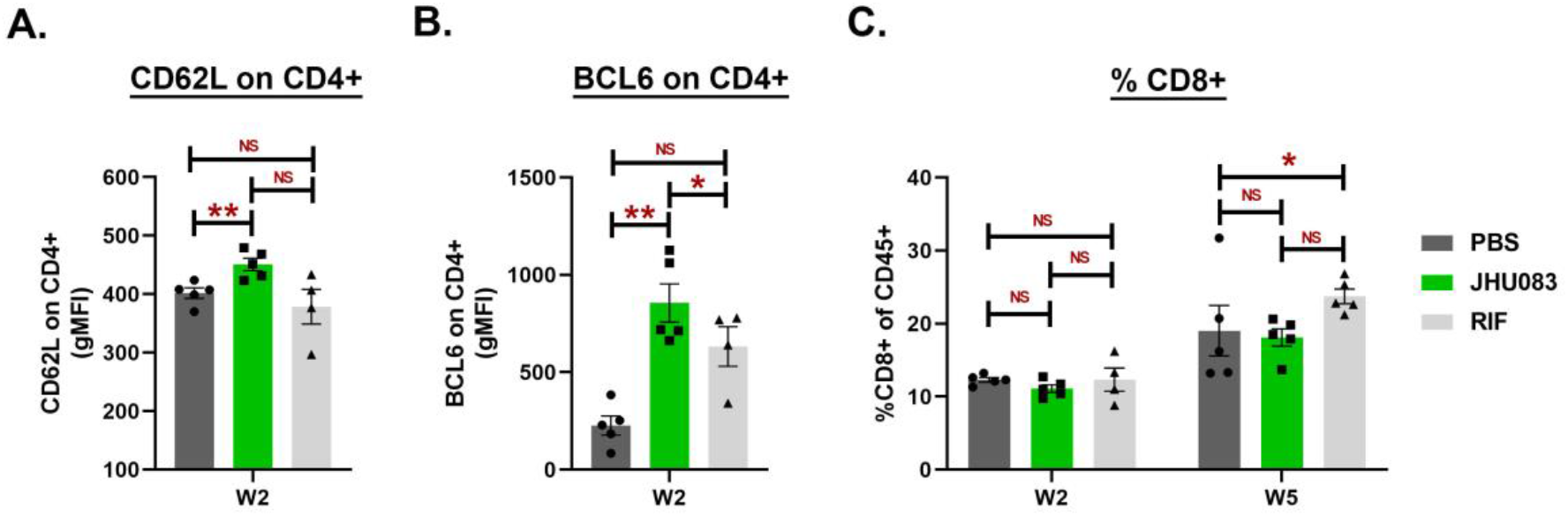
Effect of JHU083 administration upon T-cells in the lungs at weeks 2 and 5. As described in Fig 2A, *Mtb*-infected 129S2 mice were treated with JHU083 and RIF every day starting day 1 post-infection. The mice were sacrificed at week 2 and week 5, and the lungs were harvested. Single cell suspensions of the lungs from all three groups were stained with appropriate antibodies and analyzed using multicolor-flow cytometry (n=4-5). We found <u>differences</u> in the **(A)** CD62L expression upon CD4^+^ T-cells, **(B)** BCL6 expression upon CD4^+^ T-cells and, **(C)** total CD8^+^ T-cells The X-axis shows the timepoint at which the lungs were harvested for the flow cytometry analysis. Data were plotted as Mean ± SEM and are shown as the frequency of CD45^+^ population. gMFI stands for geometric mean fluorescence intensity and was used to define the expression of the individual markers upon the indicated cell types. Statistical significance was calculated using a two-tailed student t-test considering unequal distribution. *<0.05, **<0.01. The experiment was repeated two times.

**Fig S5.**
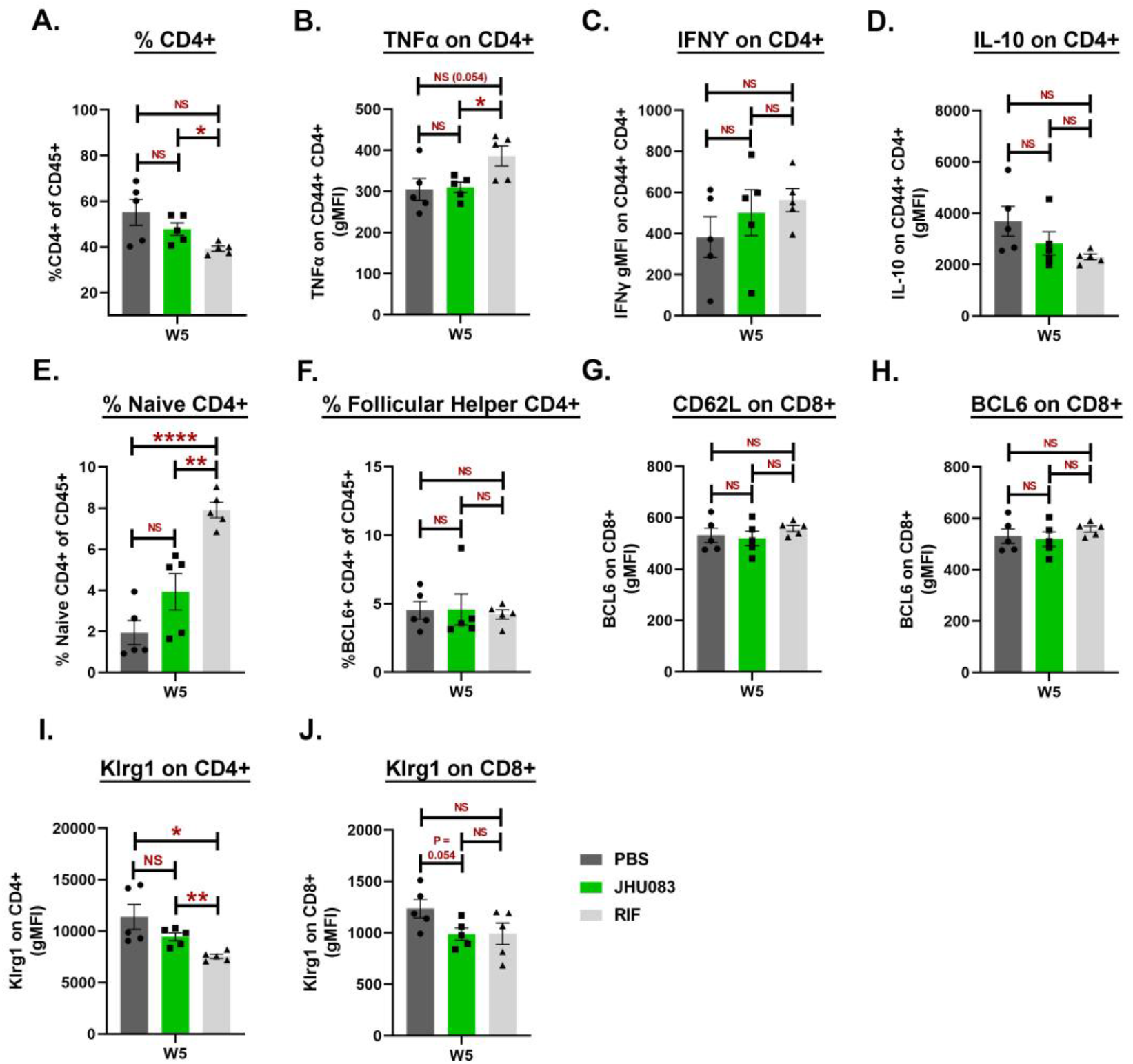
Effect of JHU083 administration upon T-cells in the lungs at week 5. As described in Fig 2A, *Mtb*-infected 129S2 mice were treated with JHU083 and RIF every day starting day 1 post-infection. The mice were sacrificed at week 2 and week 5, and the lungs were harvested. Single cell suspensions of the lungs from all three groups were stained with appropriate antibodies and analyzed using multicolor-flow cytometry (n=4-5). We found no difference in the **(A)** CD4^+^ T-cell frequency, **(B)** TNFα expression upon activated CD4^+^ T-cells, **(C)** IFNƳ expression upon activated CD4^+^ T-cells, **(D)** IL-10 expression upon activated CD4^+^ T-cells, **(E)** Naïve CD4^+^ T-cells, **(F)** Follicular helper T-cells, **(G)** CD62L expression on CD8^+^ T-cells, **(H)** BCL6 expression on CD8^+^ T-cells, **(I)** Klrg1 expression on CD4^+^ T-cells, **(J)** Klrg1 expression on CD8^+^ T-cells. The X-axis shows the timepoint at which the lungs were harvested for flow cytometry analysis. Data were plotted as Mean ± SEM and are shown as the frequency of CD45^+^ population. gMFI stands for geometric mean fluorescence intensity and was used to define the expression of the individual markers upon the indicated cell types. Statistical significance was calculated using a two-tailed student t-test considering unequal distribution. *<0.05, **<0.01, ***<0.001, ****<0.0001. The experiment was repeated two times.

**Fig S8.**
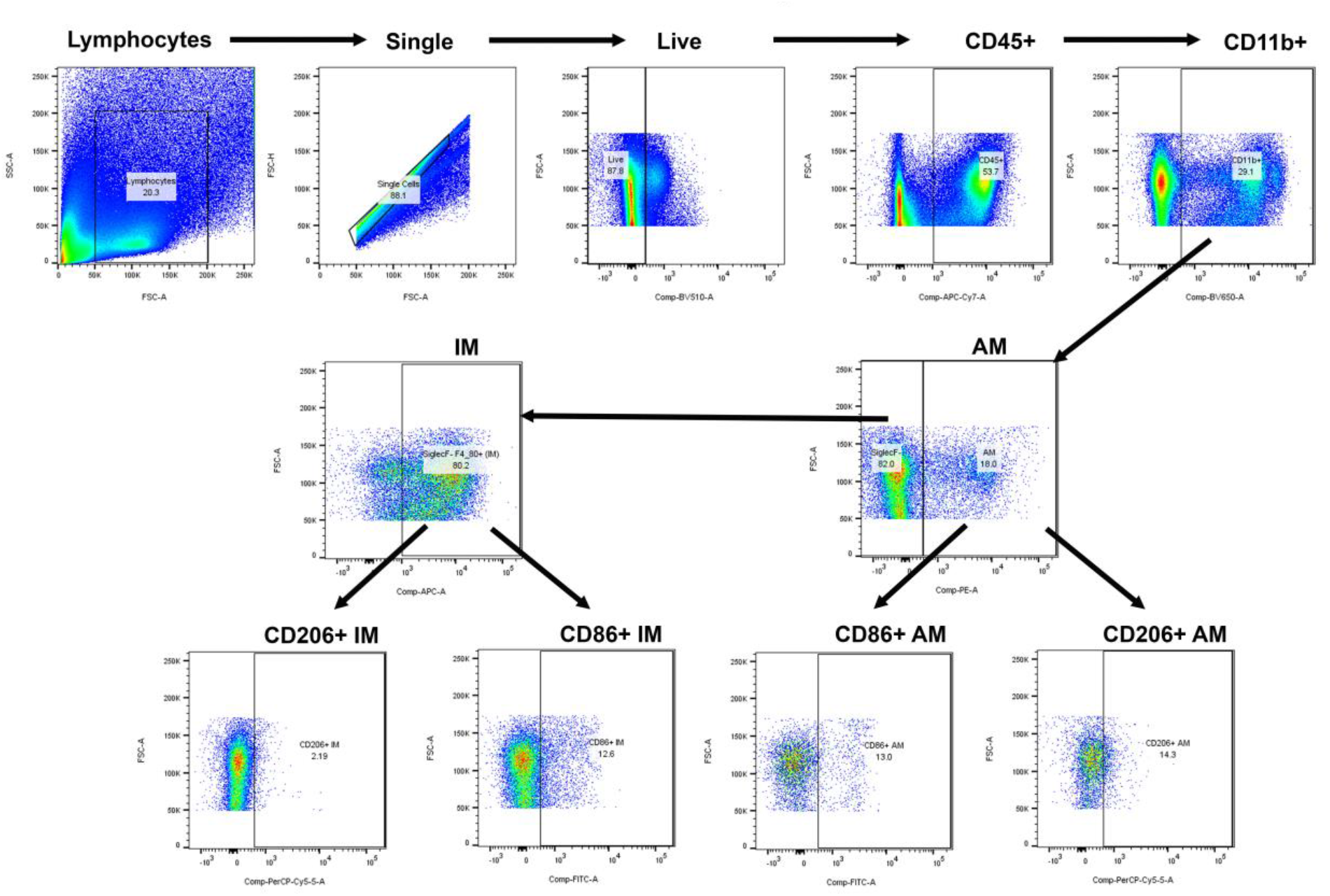
Two-dimensional gating strategies for flow cytometrical identification of myeloid cell subsets. We excluded doublets and debris and gated on single live CD45+ cells. We identified myeloid cells (CD45^+^ CD11b^+^), alveolar macrophages (AM; CD45^+^ CD11b^+^ SiglecF^+^), interstitial macrophages (IM; CD45^+^ CD11b^+^ SiglecF^-^ F4/80^+^), CD86^+^ alveolar macrophages (AM; CD45^+^ CD11b^+^ SiglecF^+^ CD86^+^), CD206^+^ alveolar macrophages (AM; CD45^+^ CD11b^+^ SiglecF^+^ CD206^+^), CD86^+^ interstitial macrophages (IM; CD45^+^ CD11b^+^ SiglecF^-^ F4/80^+^ CD86^+^), CD206^+^ interstitial macrophages (IM; CD45^+^ CD11b^+^ SiglecF^-^ F4/80^+^ CD206^+^).

**Fig S9.**
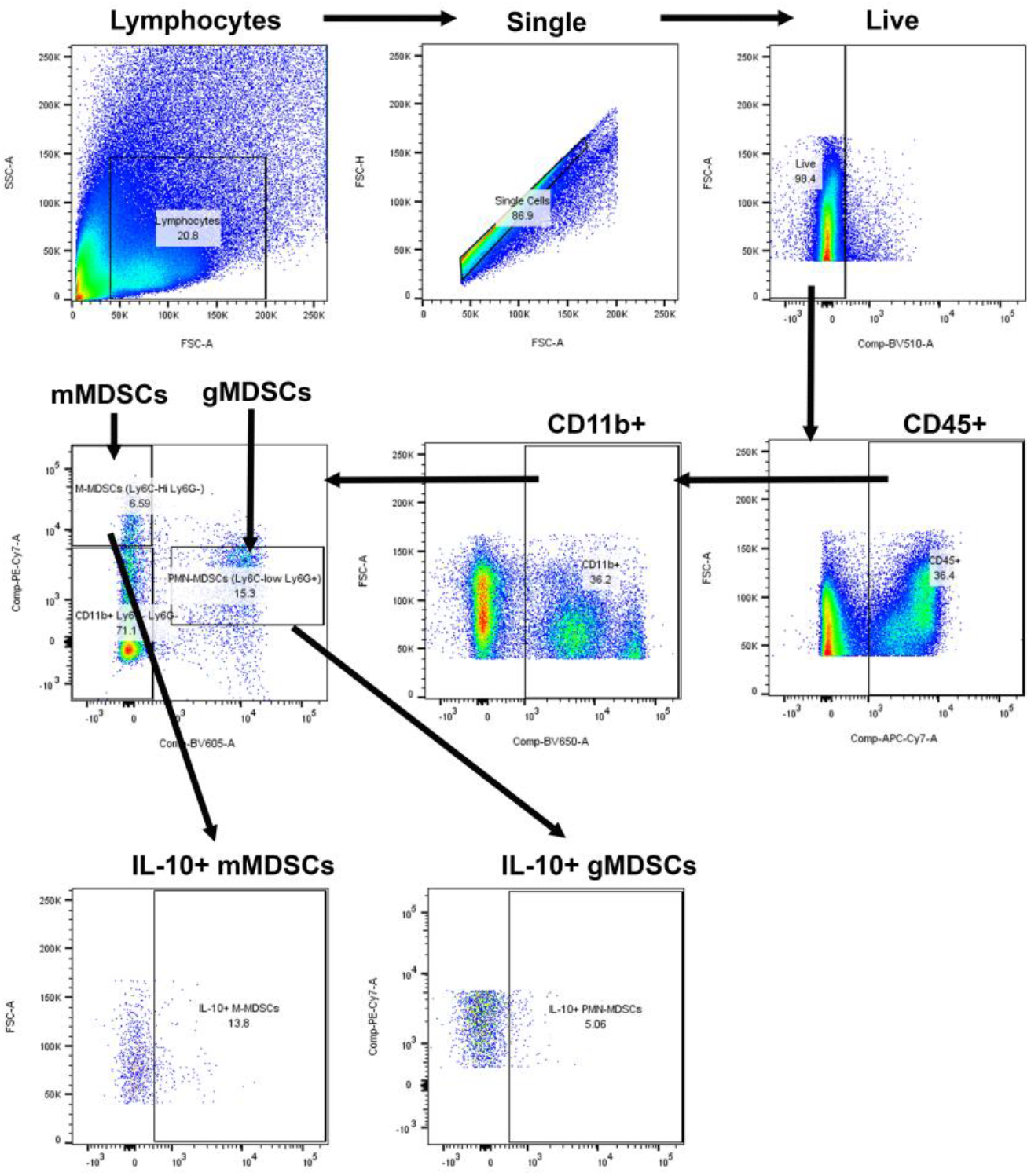
Two^-^dimensional gating strategies for flow cytometrical identification of MDSC subsets. We excluded doublets and debris and gated on single live CD45^+^ cells. We identified myeloid cells (CD45+ CD11b^+^), monocytic MDSCs (CD45^+^ CD11b^+^ Ly6G^-^ Ly6C^High^), IL-10^+^ monocytic MDSCs (CD45^+^ CD11b^+^ Ly6G^-^ Ly6C^High^, IL-10^+^), granulocytic MDSCs (CD45^+^ CD11b^+^ Ly6G^+^ Ly6C^low^), granulocytic MDSCs (CD45^+^ CD11b^+^ Ly6G^+^ Ly6C^low^, IL-10^+^).

**Fig S10.**
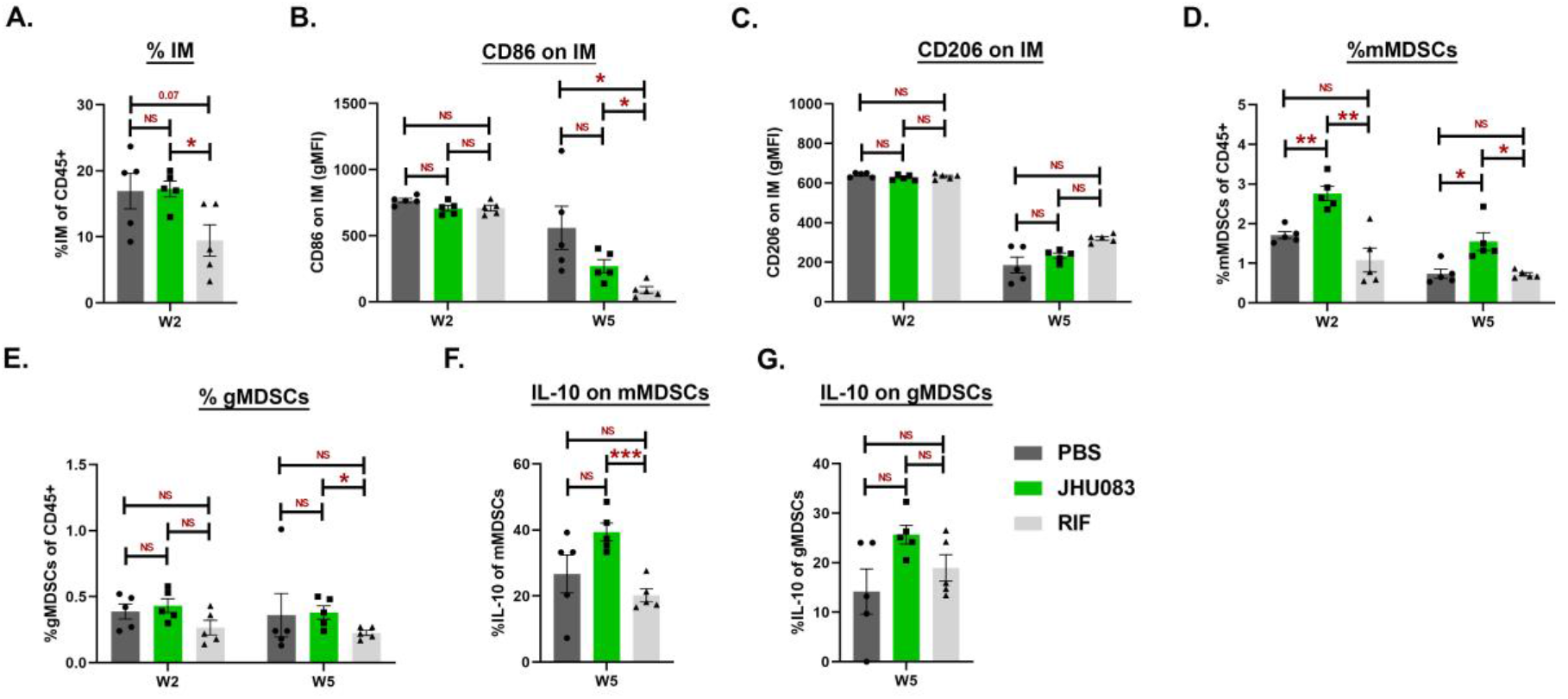
Effect of JHU083 treatment upon myeloid cell subsets. As described in Fig 2A, *Mtb*-infected 129S2 mice were treated with JHU083 and RIF every day starting day 1 post-infection. The mice were sacrificed at week 2 and week 5, and the lungs were harvested. Single cell suspension of the lungs from all three groups were stained with appropriate antibodies and analyzed using multicolor-flow cytometry (n=5). Details are provided in “Methods” section. We found no difference in the **(A)** interstitial macrophages (IM), **(B)** CD86 expression on IM, **(C)** CD206 expression on IM, **(D)** monocytic MDSC (mMDSCs), **(E)** granulocytic MDSC (gMDSCs), **(F)** IL-10 expression upon monocytic MDSCs (mMDSCs), **(G)** IL-10 expression upon granulocytic MDSCs (gMDSCs). The X-axis shows the timepoint at which the lungs were harvested for flow cytometry analysis. Data were plotted as Mean ± SEM and are shown as the frequency of CD45^+^ population. gMFI stands for geometric mean fluorescence intensity and was used to define the expression of the individual markers upon the indicated cell types. Statistical significance was calculated using a two-tailed student t-test considering unequal distribution. *<0.05, **<0.01, ***<0.001, ****<0.0001. The experiment was repeated two times.

**Fig S11.**
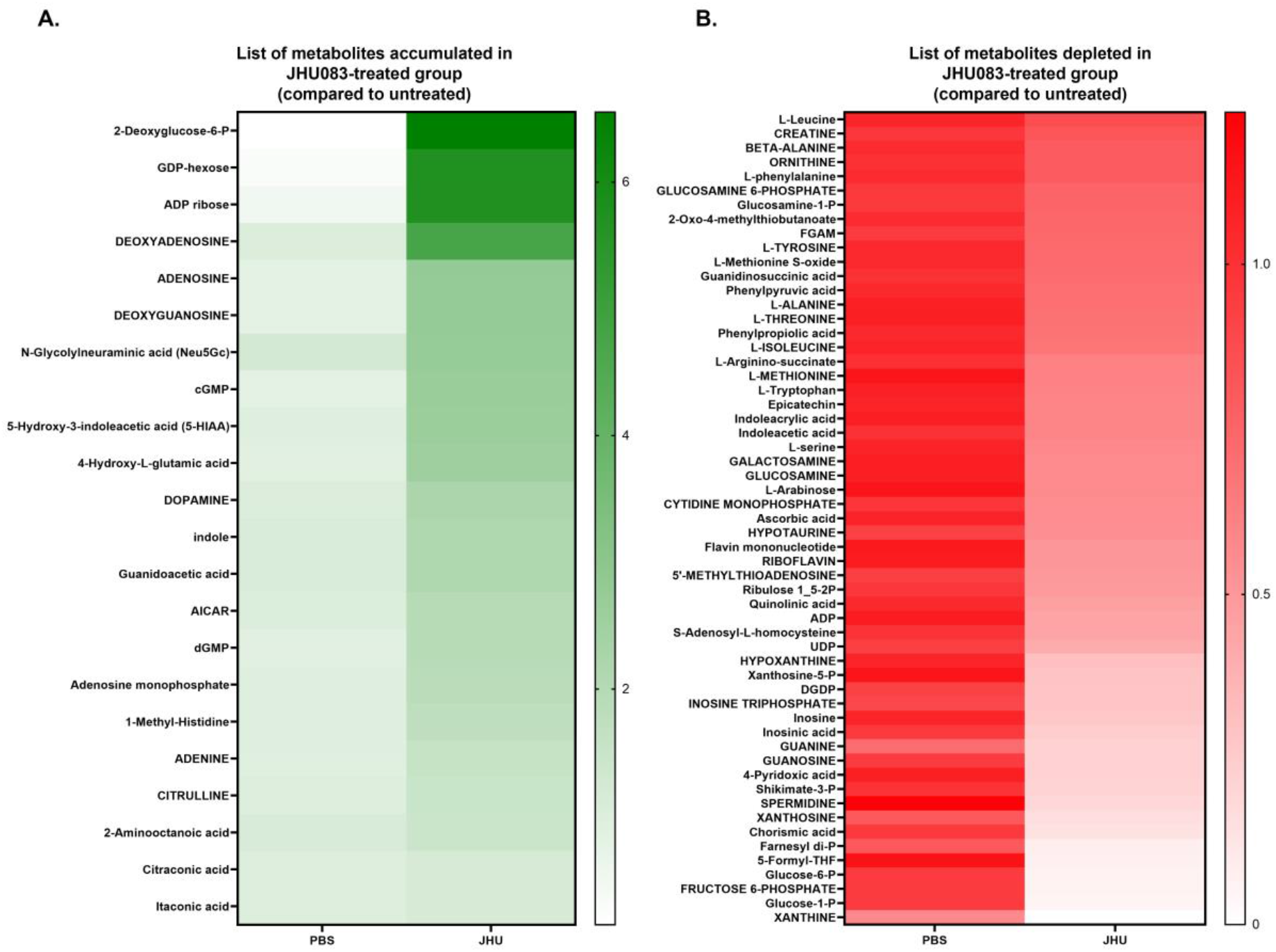
Heatmap representing the top metabolites that were either (A) enriched or (B) depleted in the lungs harvested from the animals treated with JHU083 (right-lane) compared to untreated group (left lane). The metabolites were methanol-extracted from the total lungs harvested at week 2 post infection and treatment. The metabolite abundance was normalized to the total lung tissue used for the extraction and untreated control.

**Fig S12.**
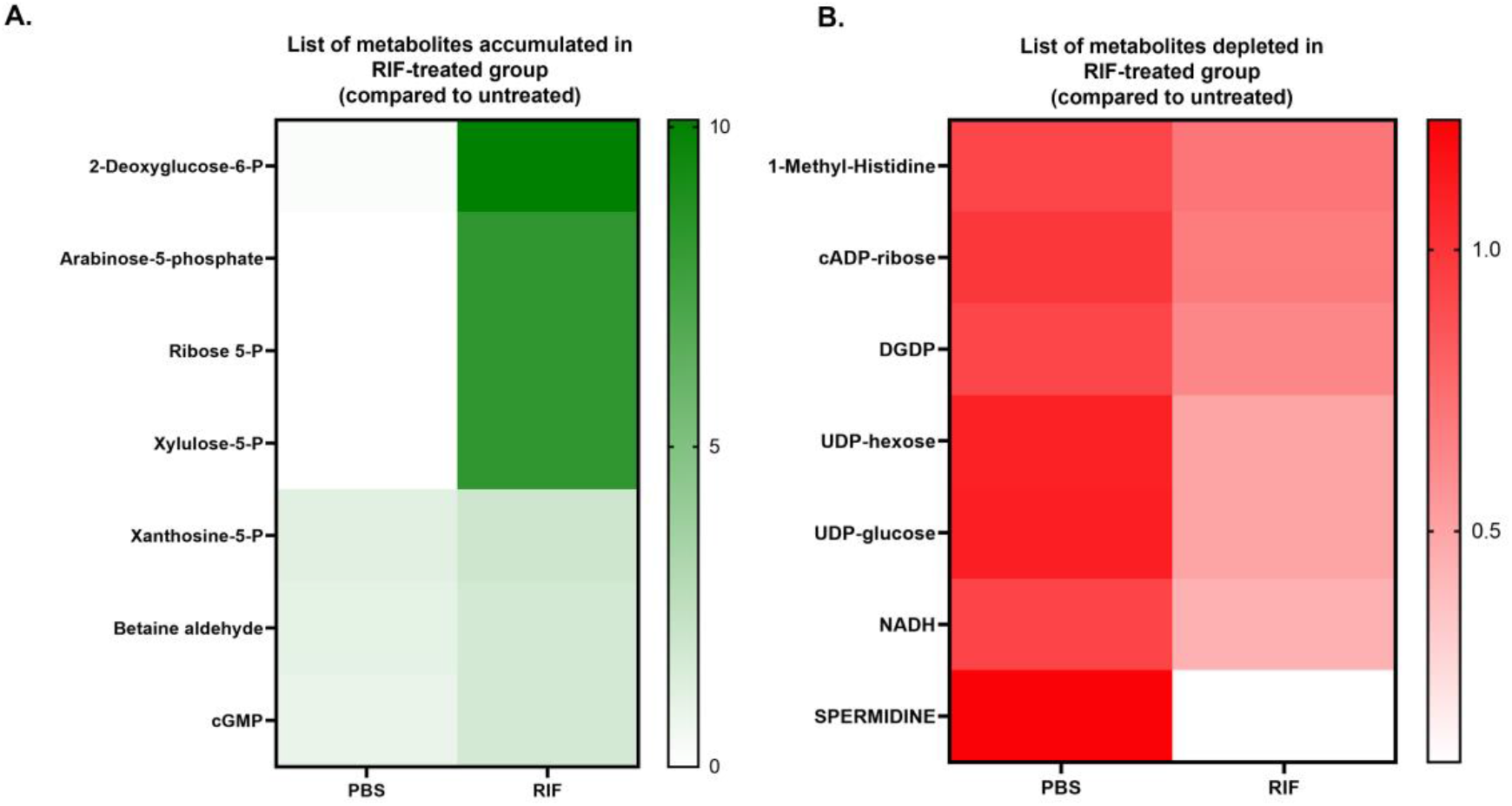
Heatmap representing the top metabolites that were either (A) enriched or (B) depleted in the lungs harvested from the animals treated with RIF (right-lane) compared to untreated group (left lane). The metabolites were methanol-extracted from the total lungs harvested at week 2 post infection and treatment. The metabolite abundance was normalized to the total lung tissue used for the extraction and untreated control.

**Fig S13.**
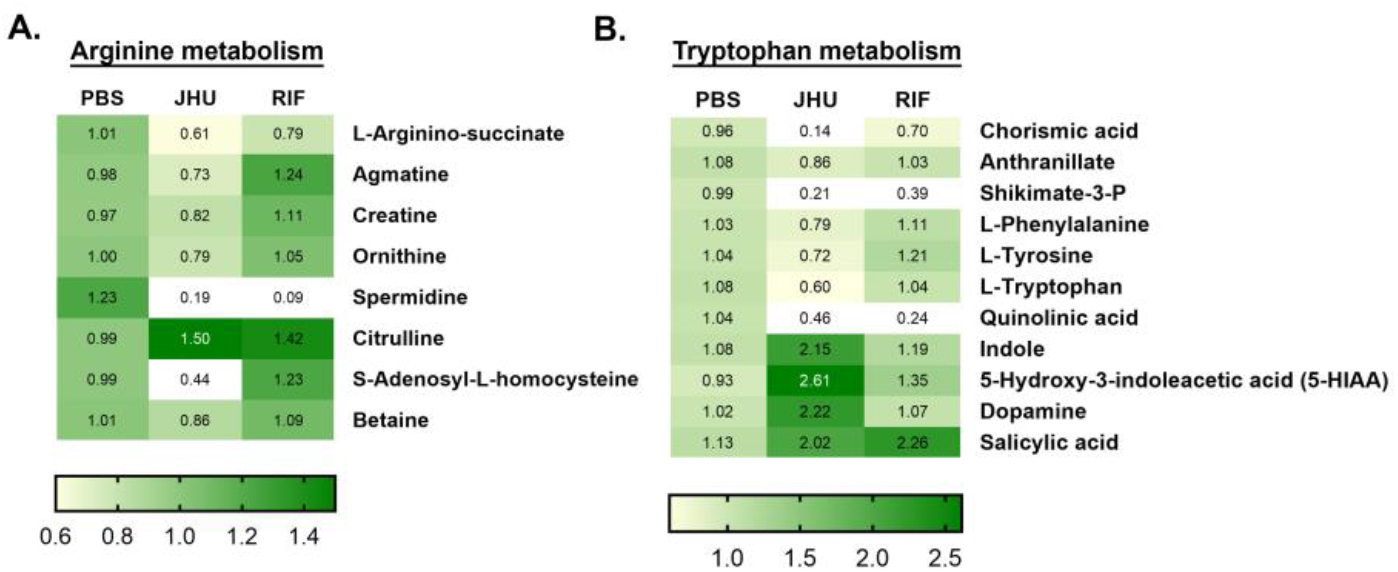
Heatmaps listing the metabolites that are affected by JHU083-treatment. As described in Fig 2A, *Mtb*-infected 129S2 mice were treated with JHU083 and RIF every day starting day 1 post-infection. Mice were sacrificed at week 2, the lungs were harvested, and total metabolites were methanol extracted as described in “Methods”. The total metabolites were normalized to the tissue weight and then to the untreated controls. We detected changes in the level of metabolites belonging to both **(A)** arginine and, **(B)** tryptophan metabolism pathways. The values in the cells correspond to the median value of the dataset. The experiment was performed once. The metabolomics data is provided as a separate excel sheet with normalized values along with the p-value calculations.

## SUPLEMENTAL DATA

## METHODS

### Alamar Blue Assay for MIC determination

The assay was performed in 96-well clear-bottom microplates as described earlier ^1,2^. Stock solutions of JHU083, DON MSO, and RIF were diluted 2-fold in 0.1 ml 7H9 broth. 0.1 ml of *Mtb* H37Rv grown to mid-exponential phase was diluted and added to each well and incubated at 37 C. After 7 days, we added 20 μl of 10X alamarBlue solution (Invitrogen) along with 12.5 μl of Tween 80 (20%) were added to the well. After overnight incubation (>16h), fluorescence intensity was measured using a fluorescence microplate reader at 544Ex/590Em nm. Minimum inhibitory concentration (MIC) was defined as the lowest drug concentration at which 90% growth inhibition was observed. RIF and no drug wells were used as positive and negative controls. We used 7H9 broth without Tween-80 addition.

### Bone-marrow-derived macrophage infection studies

BMDMs were isolated, as described earlier ^3^. Briefly, BMDMs were isolated from the femurs of 3-4 months old female C57BL/6 mice. The isolated monocytes were differentiated into macrophages in RPMI medium-Glutamax (Gibco; 61870-036) containing 10% FBS (Gibco; 16140071) and 1X antibiotic antimycotic Solution (Sigma Aldrich; A5955) and 30% (v/v) L929 conditioned media. 50,000 BMDMs were polarized using 10 ng/ml IFNϒ per well for 24 hr. The macrophages were then infected with *Mtb* H37Rv at an MOI of 2 for 4 h. The cells were washed with prewarmed media and incubated with 200 μg/ml gentamicin to eliminate the extracellular bacteria. The cells were then collected on days 1, 3, and 5, lysed with 0.25% SDS, diluted, and then plated on 7H11 selective plates. The CFU enumeration was performed on days 21-28, and the final values were plotted on a log scale of 10.

For testing the direct cytotoxic activity of DON upon macrophages, 50,000 BMDMs were plated in triplicates in 200 μl volume in a 96-well plate. The cells were then pretreated with 10 ng/ml IFNϒ per well. After 24 h, DON (1X and 10X MIC daily) and INH (32X MIC on day 1) were added. On days 1, 3, and 5 post-IFN***γ*** treatment, [3-(4,5-dimethylthiazol-2-yl)-5-(3-carboxymethoxyphenyl)-2-(4-sulfophenyl)-2H-tetrazolium, inner salt; MTS reagent (Promega) were added, and the absorbance was recorded at 490 nM with a Microplate reader (Bio-Rad, Hercules, CA). The absorbance from the wells with no cells and only media was subtracted to negate the background signal.

### Animal infection studies

Six to ten weeks old 129S2 and C3HeB/FeJ mice were procured from Charles River Laboratories (Wilmington, Massachusetts) and Jackson Laboratory (Bar Harbor, MA), respectively. 200-300 CFU of *Mtb* H37Rv strain was used for all infections. The day after infection, we randomly distributed mice into three groups: (1) PBS, (2) JHU083, and (3) RIF group. 1 mg/kg JHU083 was administered daily for the first week and three times a week afterward. 1.25 mg/kg RIF was administered daily as the positive control. All the drugs were administered orally.

For CFU enumeration, lungs were collected at weeks 2 and 5 post-infection and treatment. The lungs were then homogenized, serially diluted in PBS, and 100 μl aliquots spread on 7H11 selective plates. For the survival study, ten mice per group were kept under daily observation. C3HeB/FeJ mice were sacrificed mice at week 4 post infection and treatment.

### Lung histopathology estimation

On week 5 post-infection and treatment, left lung lobes were harvested from PBS- and JHU083-treated C3HeB/FeJ mice and stored in formalin. The formalin-fixed tissues were then sectioned and stained with Hematoxylin and Eosin (H&E). Stained sections were imaged at 40X magnification. The lesion areas were quantified using ImageScope Software (Leica Biosystems) and plotted using GradPad Prism software.

### Single-cell suspension preparation

For multicolor flow cytometry analysis, lungs and spleens were collected in 5 ml MACS Tissue storage solution (Cat: 130-100-008; Miltenyi Biotec, Gaithersburg, MD) followed by storage at 4 °C until processing. The lungs were dissected into individual lobes, and single-cell suspension was prepared using the mouse lung dissociation kit (Cat: 130-095-927; Miltenyi Biotec, Gaithersburg, MD) and GentleMACS™ Dissociator treatment following the manufacturer’s protocol. Spleens were mechanically dissociated in digestion buffer (RPMI 1640 + 10% FBS + 0.2 mg/ml collagenase D + DNAse). Both lung and spleen cells were then incubated with ACK lysis buffer at RT for 2-5 minutes to lyse the red blood cells. The cell suspension was then washed with RPMI complete media and resuspended in the appropriate volume of the same media. Trypan blue staining was performed to assess the viability of the cell suspensions.

### Multicolor flow cytometry

Lung single-cell suspensions were incubated with TruStain FcX™ (anti-mouse CD16/32) antibody (Cat: 101320; BioLegend, San Diego, CA) in eBioscience™ Flow Cytometry Staining Buffer (Cat: 00-4222-57, San Diego, CA) for 20 min at room temperature to block non-specific antibody binding. The cells were then incubated with appropriate antibody cocktails and fixation buffer (Cat: 420801; BioLegend). For intracellular staining, the cells were stained using True-Nuclear™ Transcription Factor Buffer Set (Cat: 424401; Biolegend) following the manufacturer’s protocol. Intracellular cytokine stimulation and staining was performed as described earlier ^4^. Briefly, lung cells were incubated with cell activation cocktail (Biolegend) and monensin for 4 h. The cytokine staining was performed using Cyto-Fast™ Fix/Perm Buffer Set (Cat: 426803; BioLegend) following the manufacturer’s protocol. The stained cells were stored in Cyto-Last™ Buffer (Cat: 422501; BioLegend) at 4 °C till the data acquisition. The data were acquired on BD LSRFortessa™ Cell Analyzer (BD Biosciences, San Jose, CA) and the data were analyzed using FlowJo version 10.8.1 (Tree Star). Gating strategies are provided in the supplementary **Figures S3, S4, S5, S8, S9**. Instead of beads, splenocytes were used for single-cell stain controls and for creating compensation matrices. All flow antibodies were titrated to identify the optimal concentrations for staining protocols.

### Metabolite Extraction

For LC/MS based metabolomics, lung tissues were collected from all three groups and flash-frozen in liquid nitrogen followed by storage at -80 until further processing. For the processing, 10-30 mg lungs were homogenized in 500 μL of **methanol:water (80:20, v/v) extraction solution, vortexed** and centrifuged at 14000 x *g* for 10 min at 4 °C. The supernatant fluid was stored at -80 °C for overnight to precipitate proteins. The supernatant was centrifuged at 14000 x *g* at 4 °C and filtered through 0.22 μm acetate filters (Corning, Corning, NY; Cat: 8160), lyophilized and stored at -80 °C for subsequent analysis.

### Metabolite Measurement with LC-MS

The dried metabolite extracts were resuspended in 50% acetonitrile solution. an LC-MS system consisting of an Agilent 1290 Infinity II Binary UHPLC pump and a Bruker timsTOF Pro II mass spectrometer was used for the LC-MS based metabolomics profiling.

HILIC-LC chromatographic separations were performed on the above UHPLC using a Waters XBridge BEH Amide column (2.1 × 150 mm, 1.7μm). The LC parameters were as follows: autosampler temperature, 4 °C; injection volume, 2 μl; column temperature, 40 °C; and flow rate, 0.20 ml/min. The solvents and optimized gradient conditions for LC were: Solvent A, Water with 0.1% formic acid; Solvent B, Acetonitrile with 0.1% formic acid; A non-linear gradient from 99% B to 45% B in 25 minutes with 5min of post-run time.

MS spectra were collected using a timsTOF Pro II mass spectrometer (Bruker Daltonics) equipped with IonBooster ESI source. The mass spectrometer was operated in negative mode with auto MS/MS method. The optimized operation parameters were End Plate Offset, 400V; Capillary Voltage,1000 V; Charging Voltage, 300 V; Nebulizer Pressure, 4.1 bar; Dry Gas 3 L/min; Dry Gas Temperature, 200 °C; Vaporizer Temperature, 350 °C; The mass scan range, 70-1100 m/z; Scan rate, 12 Hz. Spectra were internally mass calibrated at the beginning of every sample run by infusion of a small fragment of reference mass solution using a syringe pump connected to the sprayer feeding into the ESI source. Data were acquired with Compass HyStar 5.1 acquisition software and processed with TASQ 2022. The analyte database used for metabolite identification was developed in house with retention times based on the HILIC method. We note that our methods do not distinguish some metabolites with the same formula and very similar structures, hence care should be taken in interpretation of such data.

---

**SI Table 1.**
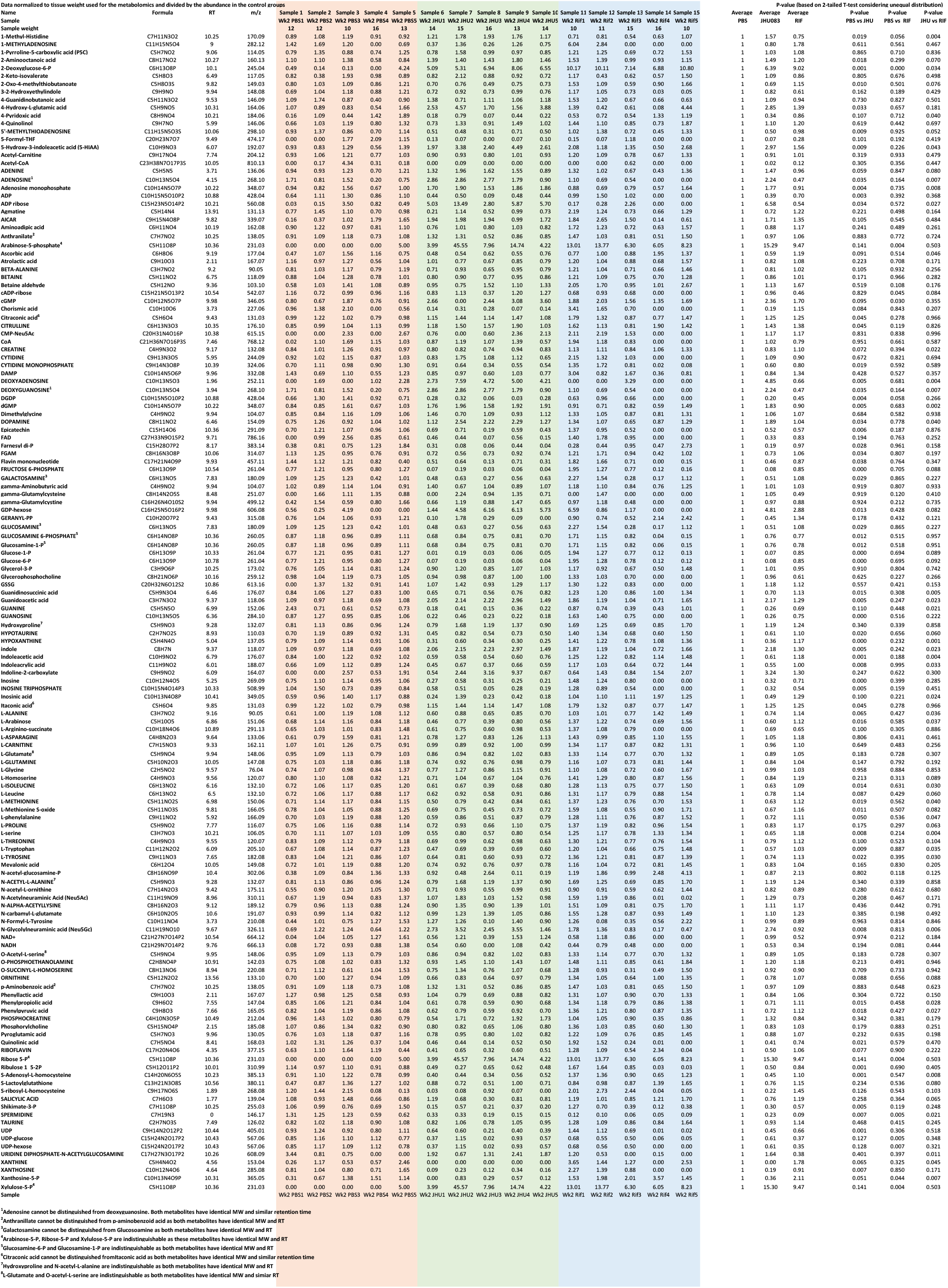
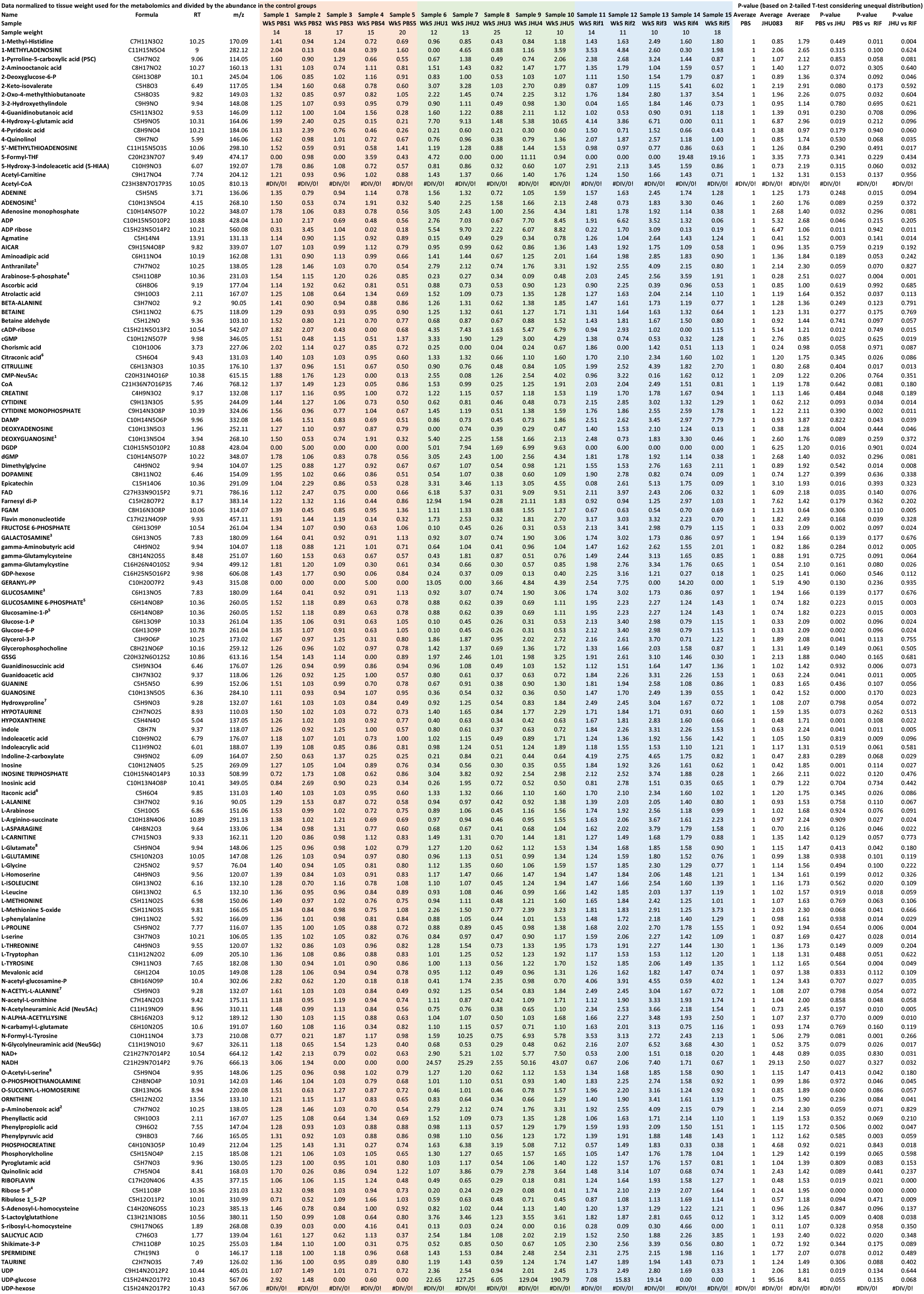
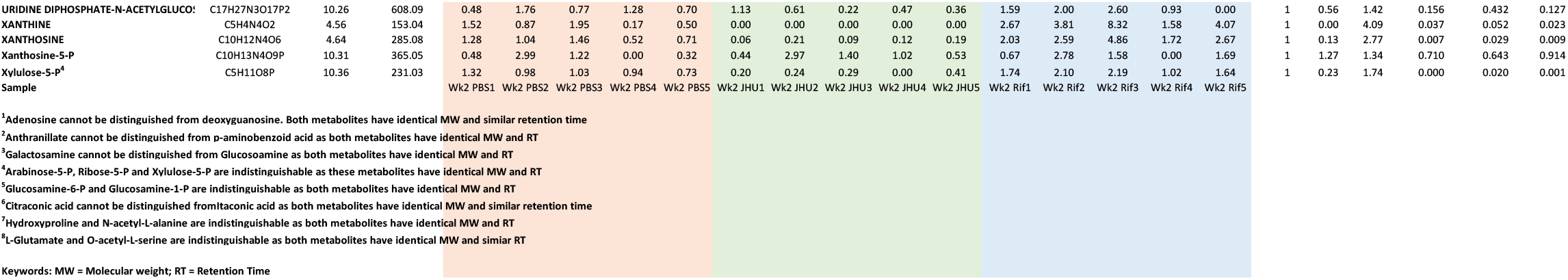
List of all the metabolites identified in the metabolomics (Week 2) List of all the metabolites identified in the metabolomics (Week 5)

